# Inhibition of Tgfβ signaling enables durable ventricular pacing by TBX18 gene transfer

**DOI:** 10.1101/2022.06.02.493572

**Authors:** Jinqi Fan, Nam Kyun Kim, Natasha Fernandez, Tae Yun Kim, Jun Li, David Wolfson, Hee Cheol Cho

## Abstract

Implantable cardiac pacemaker devices are generally effective for patients with symptomatic bradyarrhythmia. However, device-dependent cardiac pacing is far from ideal and often inadequate, particularly for pediatric patients who need to go through invasive revision of the indwelling hardware. Biological pacemakers have been proposed as device-free alternatives to the current treatment, but sustained, unwavering biological pacing beyond days after the biologic delivery has not been demonstrated. We have previously demonstrated that re-expression of an embryonic transcription factor, TBX18, could reprogram ventricular cardiomyocytes into induced pacemaker myocytes (iPMs). Here, we report that exogenous expression of TBX18 *per se* leads to severe fibrosis *in situ*, impairing the iPMs’ ability to pace together. Acute fibrosis is accompanied with proliferation and activation of cardiac fibroblasts via Tgfβ-Smad2/3 pathway. Small molecule inhibition of Tgfβ signaling mitigated the interstitial remodeling, independent from TBX18-induced iPM reprogramming at the single-cell level. Direct and focal gene transfer of TBX18 into the left ventricular myocardium created ventricular pacing in a rat model of chronic atrioventricular block, but such activity began to wane in a week. In contrast, a combination therapy consisting of TBX18 gene transfer and Tgfβ inhibition enabled sustained biological pacing beyond the four-week study period. Our data demonstrate that inhibition of Tgfβ signaling suffices to achieve durable cardiac pacing by TBX18-induced biological pacemakers.

## INTRODUCTION

An abnormally slow heart rhythm, bradyarrhythmia, can be a life-threatening condition. The only treatment is surgical implantation of an electronic pacemaker. Although the pacemakers work well, complications inherent to the indwelling devices include generator failure, lead fracture, infections, and/or unintended disabling by commercial devices (*1, 2*). These problems continue to pose challenges to patients and caregivers, particularly for pediatric and adult patients with congenital heart diseases. Device pacing is far from ideal and often inadequate for newborns and infants with symptomatic bradyarrhythmia, requiring surgical repositioning and/or replacement of the hardware (*3*).

Biological alternatives to implantable devices have been proposed at the proof-of-concept level (*4, 5*). The concept has not been reduced to a viable clinical treatment, partly because its efficacy has not been demonstrated to sustain beyond days. We have previously demonstrated that re-expression of a human embryonic transcription factor, TBX18, reprogrammed postnatal ventricular cardiomyocytes into induced pacemaker myocytes (iPMs), recapitulating the hallmark features of sinoatrial node (SAN) pacemaker cells (*6*). Focal delivery of *TBX18* to the ventricular myocardium provided proof-of-principle for reprogramming-based cardiac pacing in both small and large animal models of bradyarrhythmias (*6–8*). However, *TBX18*-induced cardiac pacing was transient, and began to wane one week after the gene transfer (*7, 8*).

Tbx18 is required for embryonic development of the SAN (*9*), but its expression is not exclusive to the SAN. The proepicardium of the developing heart is marked by progenitors that express Tbx18 and Wt1, which differentiate into epicardial cells and fibroblasts (*10, 11*). Reactivation of Tbx18 in the postnatal epicardium appears to be correlated with pathological fibrosis in the ischemic or hypertensive heart (*12, 13*). The SAN of the healthy heart is characteristically fibrotic, with estimates of >50% of the SAN volume to be occupied by nonmyocytes (*14*). Noting that adenoviral somatic gene transfer does not discriminate between cardiomyocytes and nonmyocytes *in vivo*, we set out to understand the consequences of *TBX18* expression in the context of extracellular matrix remodeling, fibrosis, and cell-cell coupling. We hypothesized that re-expression of exogenous *TBX18* in the chamber myocardium not only creates de novo pacemaker cells but also triggers fibrosis, compromising TBX18-iPMs’ ability to pace in synchrony. Our data indicate that exogenous expression of *TBX18* triggers proliferation and activation of cardiac fibroblasts (FBs), which is mitigated by inhibition of Tgfβ signaling. Transient inhibition of Tgfβ signaling sufficed to rescue the loss of automaticity in TBX18-iPMs *in vitro* and generated unwavering ventricular pacing *in vivo* for more than four weeks in a rat model of complete and chronic heart block.

## RESULTS

### TBX18 re-expression elicits significant fibrosis in situ

In view of the potential association of pathologic fibrosis with reactivation of embryonic TBX18 in human disease, we tried to answer whether TBX18 re-expression will activate fibroblasts and elicit overt fibrosis. An adenoviral vector expressing human TBX18 and a ZsGreen reporter in a bicistronic manner (hereon Adv-TBX18) was injected directly into the left ventricular (LV) apex of the adult rat heart. Masson’s trichrome and picrosirius red staining revealed that the degree of fibrosis and collagen deposits were significantly higher at the injection site of Adv-TBX18 injected heart compared to Adv-GFP injected myocardium at days 7 (d7) and 14 after the gene transfer (Fig. 1A & 1C). The fibrotic region appeared to intensify from d7 to d14 after TBX18 gene delivery, coinciding with the waning of TBX18-induced ventricular pacing in previous studies (*7, 8*). Significantly more vimentin^+^ nonmyocytes were observed at the site of TBX18-injected hearts than in control animals injected with Adv-GFP at the same dose at d28 (Fig. 1B & 1C). To test for profibrotic effects of TBX18 without potential complications from needle injury, neonatal rat ventricular myocytes (NRVMs) were harvested and transduced with either Adv-TBX18 or Adv-GFP *in vitro*. The NRVMs include a minor population of FBs carried over during cell isolation step (fig. S1). Two weeks after the gene transfer, TBX18-NRVMs showed significantly higher vimentin^+^ FBs and alpha smooth muscle actin (αSMA) ^+^ myofibroblasts (myoFBs) compared to GFP-NRVMs (1.6-fold and 1.9-fold, respectively, n=5 biological replicates in each group, *P*<0.05). Similarly, collagen1^+^, αSA^−^ nonmyocytes were more numerous in TBX18-NRVMs than in GFP-NRVM monolayers (Fig. 1D, n=5 biological replicates in each group, *P<*0.05). Thus, re-expression of TBX18 elicits substantial fibrosis *in situ* accompanied by proliferation and activation of fibroblasts.

**Fig. 1.**
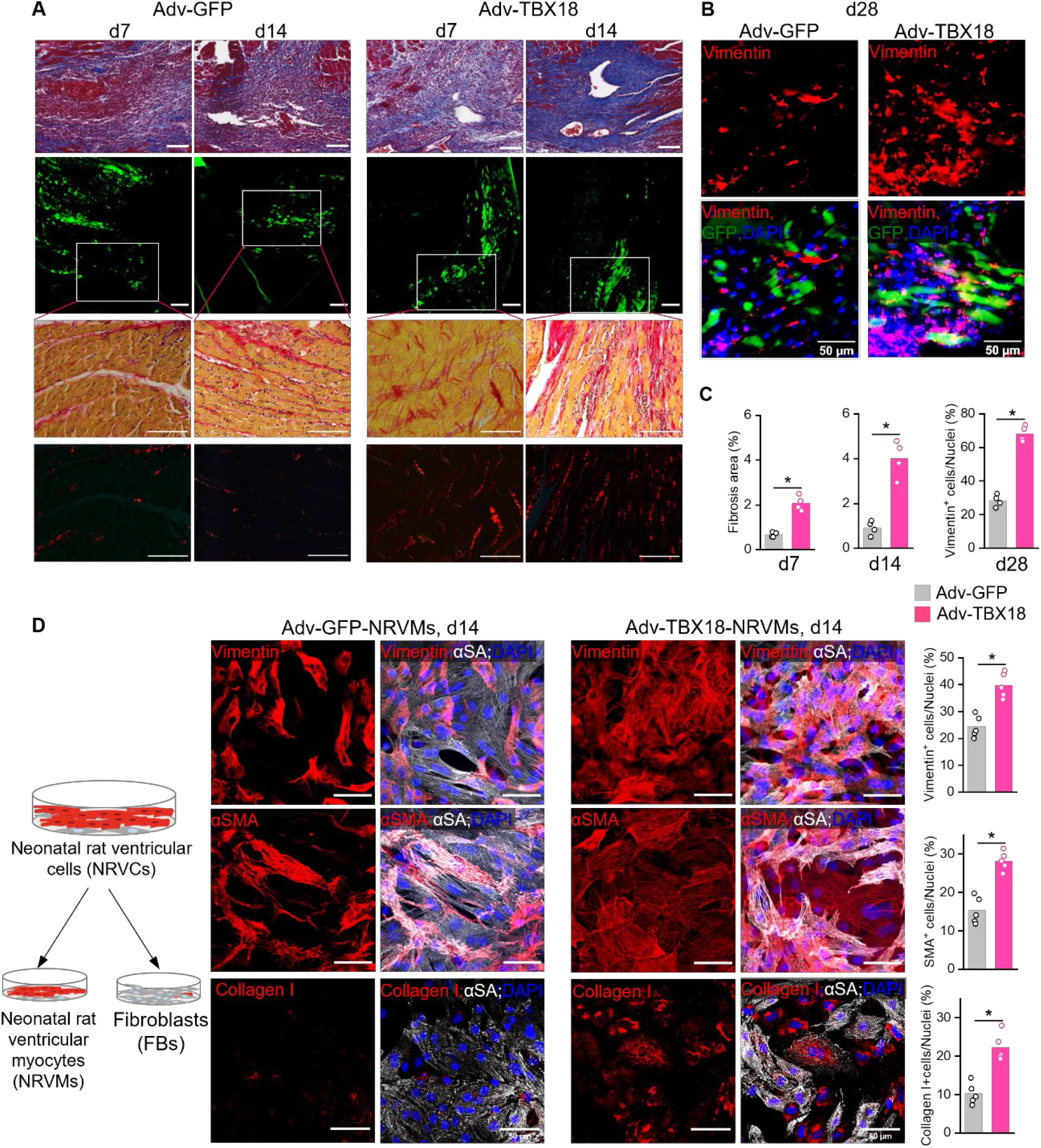
TBX18 elicits intense fibrosis. (**A**) Masson’s trichrome (top row) and Picrosirius red staining (bottom two rows) images of GFP or Tbx18-injected adult rat heart at d7 and d14 after gene delivery *in vivo*. Green fluorescence images (second row) indicate GFP or TBX18-positive myocardium in parallel sections of Picrosirius red-stained sections. Scale bar=200 µm for green channel and 100 µm for others. (**B**) Immunofluorescence images of vimentin^+^ cells (upper panel) at the site of transgene injection (GFP or Zsgreen positive cells, lower panel). (**C**) Left and middle graphs: quantification of fibrotic area analyzed from the polarized light imaging of Picrosirius staining (bottom row in A). Right: quantification of vimentin^+^ cells in B (n=4 animals in each group, *P<*0.05). (**D**) Immunostaining images of NRVMs against αSMA, vimentin and collagen I at d14 after gene transfer. Scale bar=50 µm (**B, D)**. n=5 wells per group, \**P<*0.001, Student’s t-test.

In order to understand the gene expression profile of acute fibrosis elicited by TBX18, we performed bulk RNA-seq on NRVMs transduced with either Adv-TBX18 or Adv-GFP at d4 after gene transfer. Analysis of differentially expressed genes (DEG) illustrated that among the 17,603 total gene transcripts, 934 genes were upregulated and 463 were downregulated (fig. S2A). Gene ontology (GO) enrichment analysis indicated that TBX18 re-expression activated gene regulatory networks for extracellular matrix (ECM), fibrosis and inflammation (fig. S2B). Pathway analysis showed that the upregulated DEGs were significantly enriched in Tgfβ signaling, extracellular matrix organization and cytokine production (fig. S2C). These results demonstrate that TBX18 re-expression leads to acute fibrosis and ECM remodeling.

### Tgfβ signaling mediates the fibrosis elicited by TBX18

To gain a cell-type-specific understanding of TBX18-elicited fibrosis, we performed single-cell RNA sequencing (scRNA-seq) at d3 after gene transfer into neonatal rat ventricular cells (NRVCs) from the preplating step, which was deliberately mixed with NRVM pool and nonmyocytes to better mimic myocardial cell constituents (Fig. 1D). NRVC Upon QA/QC filtering, 13,437 cells met the quality thresholds for subsequent analysis. We focused on two cell types: cardiomyocytes (CMs, *Actn2*^*high*^, *Tnni3*^high^), total FBs (*Pdgfrb*^*+*^ and *Tcf21*^+^) with 2 subpopulations, including myoFBs (*Pdgfrb*^*+*^, *Tcf21*^*+*^, *Acta2*^*high*^) and quiescent FBs (*Pdgfrb*^*+*^, *Tcf21*^*+*^, *Acta2*^*low*^) (Fig. 2A and 2B). Pathway analysis further revealed that upregulated DEGs by TBX18 in CMs and FBs are involved in fibroblast activation and ECM remodeling (Fig. 2C and 2D). Among those, Tgfβ signaling, which is one of the major molecular mediators of fibrosis in the heart (*15*), figured prominently in TBX18-transduced CMs and FBs (Fig. 2D). This was evidenced by upregulated expression of Tgfβ ligands and profibrotic genes such as *Tgfb2, Tgfb3*, and *Acta2* in TBX18-NRVCs (Fig. 2E, fig. S3). Accordingly, higher proportion of activated myoFBs (39.3% vs. 25.4%, *P*<0.001) and lower proportion of quiescent FBs were observed in TBX18-NRVCs compared to control (Fig. 2F).

**Fig. 2.**
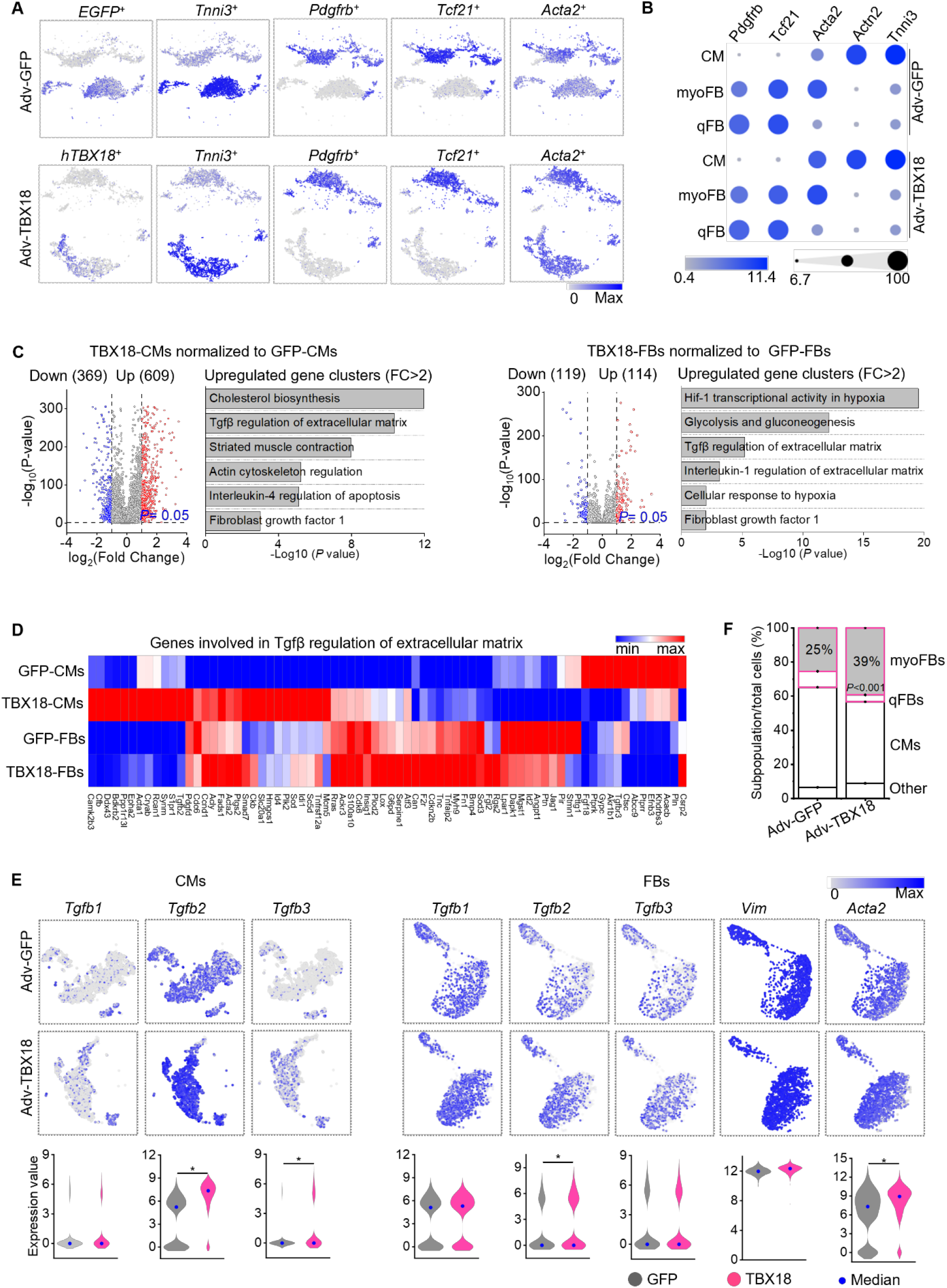
TBX18 activates myoFBs via Tgfβ signaling. (**A**) Single cell RNA-seq analysis of NRVCs transduced with either Adv-TBX18 or GFP at d3 after gene transfer. Data are presented as t-SNE plots, pseudo-spacing 13,437 cells (total, including both GFP and TBX18 groups) by nonlinear transformation of gene expression similarity. (**B**) Bubble plots indicating gene expression level (color intensity) and the percentage of positive cells (circle size) for three cell groups: cardiomyocytes (CMs), quiescent fibroblasts (qFBs), myofibroblasts (myoFBs). (**C**) Left two panels: Volcano plot of DEGs and pathway analysis of upregulated genes in TBX18-CMs normalized to GFP-CMs with threshold of |FC|>2 and *P<*0.05. Right two panels: similar to the left two panels, but with TBX18-FBs normalized to GFP-FBs. (**D**) A heat map of genes involved in Tgfβ regulation of extracellular matrix shows upregulation of fibroblast activation-related genes in TBX18-CMs and TBX18-FBs at d3. (**E**) t-SNE plots (top and middle rows) and Violin plots of gene expression (bottom) exhibit significantly upregulated expression of Tgfβ-related genes in TBX18-CMs (left three columns) and TBX18-FBs (right five columns). \**P<*0.001 compared to Ad-GFP and |FC|>1.5, Partek GSA algorithm. (**F**) TBX18-NRVCs contained higher proportion of myoFBs compared to GFP-NRVCs. *P<*0.001, Chi-square test.

To test the causative role of Tgfβ signaling in triggering fibrosis upon TBX18 re-expression, monolayers of NRVCs were treated with a small molecule inhibitor of type I Tgfβ receptors, A83-01 (*16–19*). Treatment with A83-01 significantly decreased the population of myoFBs from 68.0% in TBX18-NRVCs to 49.5% of total cell population at d14 (*P*<0.001, Fig. 3A). Heat map analysis of all genes indicated that TBX18-CMs showed upregulation of fibroblast-related genes, which was mitigated with A83-01 treatment (Fig. 3B). The upregulated DEGs in TBX18-FBs pointed to ECM remodeling as well as inflammatory response involving Tgfβ and other cytokines (Fig. 3C, right). Again, treatment with A83-01 abrogated the rise of these pathways (Fig. 3C, left; fig, S4). Profibrotic genes such as *Tgfb2, Tgfb3, Acta2*, and inflammatory genes were upregulated in FBs transduced with Adv-TBX18, which was mitigated by A83-01 treatment (Fig. 3D, fig. S4). The upregulated gene clusters in TBX18-FBs were also upregulated in TBX18-CMs, particularly, interferon signaling, interleukin-4 regulation of apoptosis (fig. S5 and S6), which may lead to the loss of TBX18-CMs over time. The inflammatory and profibrotic pathways were suppressed by treatment with A83-01 (fig. S5 & S6) and may explain the higher population of TBX18-CMs with A83-01 at d14 (Fig. 3A).

**Fig. 3.**
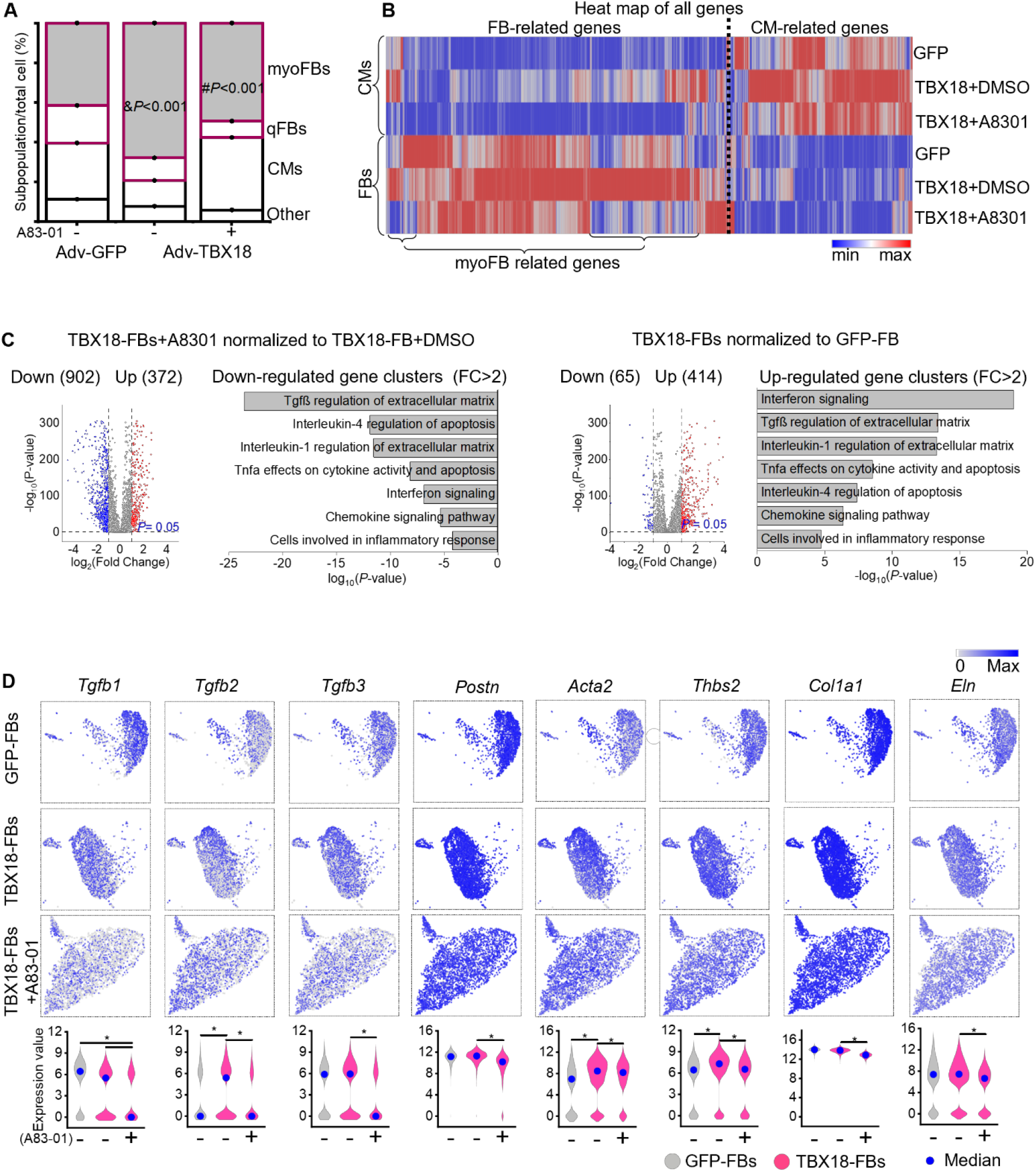
Tgfβ inhibition mitigates myoFB activation by TBX18 at d14. Data from single cell RNA-seq analysis of NRVCs transduced with Adv-GFP, Adv-TBX18 or Adv-TBX18 with A83-01 at d14 after gene transfer. (**A**) Proportion of myoFBs is higher in TBX18-NRVCs than in GFP-NRVCs at d14, which is countered by A83-01 treatment. ^&^*P<*0.001 between TBX18-NRVCs and GFP-NRVCs, ^#^*P<*0.001 between TBX18-NRVCs with and without A83-01. Chi-square test. (**B**) Hierarchical clustering of DEGs demonstrates increased expression of fibroblast-related genes in TBX18-CMs compared to GFP-CMs, which was suppressed by A83-01. Similarly, FB and myoFB-related gene expression was higher in TBX18-FBs than in GFP-FBs, which was inhibited by A83-01. (**C**) Left two panels: Volcano plot of DEGs and pathway analysis of downregulated genes in TBX18-FBs treated with A83-01 normalized to TBX18-FBs with threshold of |FC|>2 and *P<*0.05. Right two panels: similar to the left two panels, but with TBX18-FBs normalized to GFP-FBs. (**D**) t-SNE plots (top three rows) and Violin plots of gene expression (bottom) exhibit significantly upregulated gene expression of Tgfβ signaling in TBX18-FBs compared to GFP-FBs and TBX18-FBs with A83-01. \**P<*0.001 and |FC|>1.5, Partek GSA algorithm.

We validated the scRNA-seq data by quantifying the transcript levels of Tgfβ ligands and receptors. *Tgfb1, Tgfb2 and Tgfb3* as well as *Tgfbr2* were expressed higher in TBX18-NRVMs compared to control (fig. S7A). At the protein level, the concentration of secreted Tgfβ1 ligand was 70±16% higher (n=17, *P<*0.05) in the conditioned media from TBX18-NRVMs compared to that from GFP-NRVMs at d5 after gene transfer (fig. S7B). Inhibition of Tgfβ signaling with A83-01 markedly reduced the number of vimentin^+^ and αSMA^+^ cells in TBX18-NRVMs compared to untreated TBX18-NRVMs at d7 (fig. S8A). In addition, lower expression of fibrosis-related genes and proteins were observed when TBX18-NRVMs were incubated with A83-01 (fig. S8B & 8C). Phosphorylation of Smad2/3 is a key mediator of canonical Tgfβ signaling cascade (*20–22*). Phosphorylated Smad2 (pSmad2) protein level was significantly higher in TBX18-NRVMs compared to control while the total Smad2/3 protein levels were unchanged. Treatment with A83-01 completely suppressed phosphorylation of Smad2/3 in TBX18-NRVMs to the level observed in control (fig. S8B, right). Thus, the data support that TBX18 elicits myoFBs activation through Tgfβ/pSmad2/3 signaling.

We asked whether activation of Tgfβ signaling by TBX18 in cardiomyocytes could trigger fibrosis in a paracrine manner. A transwell system was employed to culture GFP or TBX18-transduced NRVMs monolayers on the semipermeable inserts, separating the NRVMs from non-transduced primary FBs seeded on the bottom (fig. S8C, bottom). Transcript levels of *Tgfβ* signaling and ECM genes, such as *Col1a1, Col3a1, Eln* and *Tgfb1*, were higher in FBs cultured in the same media with TBX18-NRVMs inserts than those with GFP-NRVMs inserts. Addition of A83-01 in the culture media abrogated the increase in Tgfβ and ECM genes (fig. S8C, bottom). Interestingly, gene transfer efficiency was significantly lower in FBs compared to CMs upon transduction of NRVCs with Adv-TBX18 or Adv-GFP. Among CMs, the transduction efficiency was 28.6% with Adv-TBX18 and 56.3% with Adv-GFP at d3. Among FBs, the gene transfer efficiency was only 2.5% and 7.1% with Adv-TBX18 or Adv-GFP (fig. S9, left), respectively. These data indicate that TBX18 activates FBs indirectly via paracrine Tgfβ signaling of cardiomyocytes.

### TBX18-induced pacemaker cell reprogramming is unaffected by Tgfβ inhibition

We asked whether the Tgfβ signaling-mediated fibrosis impacts TBX18-induced nodal cell reprogramming. We quantified the number of de novo iPMs based on: *Tnni3*^*high*^, *Hcn4*^*+*^, *Nkx2-5*^*low*^, *Gja1*^*low*^ through our scRNA-seq analysis and demonstrated the proportions of *Hcn4*^*+*^ iPMs was similar in TBX18-CMs regardless of Tgfβ inhibition but was higher compared to control (Fig. 4A & 4B), which was confirmed by the immunostaining of Hcn4^+^ in TBX18-NRVMs with or without Tgfβ inhibition (Fig. 4C) (6). Transcript levels of ion channel genes that are enriched in nodal pacemaker myocytes (*Hcn4*) or the chamber cardiomyocytes (*Kcnj2 and Scn5a*) were regulated similarly in TBX18-NRVMs with and without Tgfβ inhibition (Fig. 4D). We previously observed that gene transfer of TBX18 *in vivo* changes the morphology of rod-shaped ventricular myocytes to spindle-shaped nodal-like myocytes (*6*), similar to the morphology of native SAN myocytes (*23*). Here, we examined the cell morphology of freshly isolated ventricular cardiomyocytes at three weeks after gene transfer into the adult rat ventricular myocardium with or without A83-01 treatment for the first 7 days with an osmotic pump. Single TBX18^+^ (zsGreen^*+*^) cardiomyocytes from animals treated with either A83-01 or vehicle-only exhibited long and tapering ends akin to the morphology of the nodal pacemaker cells (Fig. 4E). The width-to-length ratio of TBX18-transduced cardiomyocytes was higher compared to that of control animals injected with GFP only but was indistinguishable between TBX18-injected animals treated with A83-01 or vehicle only (Fig. 4E). Together, the findings indicate that inhibition of Tgfβ signaling does not impact TBX18-induced reprogramming of chamber cardiomyocytes to iPMs.

**Fig. 4.**
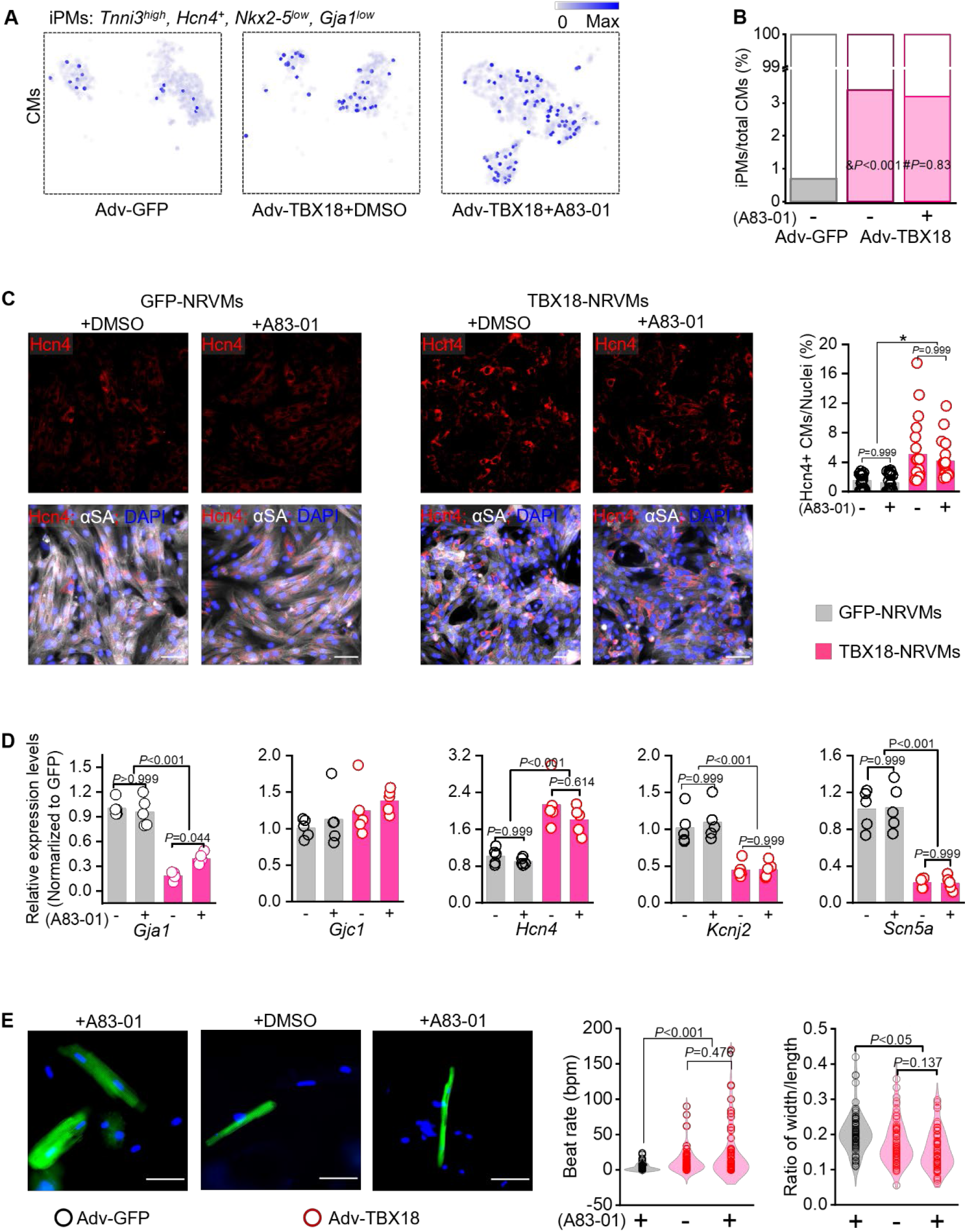
Tgfβ pathway does not interfere with TBX18-iPM reprogramming. (**A**) t-SNE plot identifying iPMs defined by Tnni3^high^, Hcn4^+^, Nkx2-5^low^ and Gja1^low^. **(B)** The proportion of iPMs among total CMs is similar between TBX18-treated NRVCs with or without A8301 (#*P*=0.83). &*P*<0.001 compared to GFP-CM, Chi-square test. (**C**) Hcn4 immunostaining shows higher ratio of Hcn4^+^ cells in TBX18-NRVMs with and without A8301 compared to control. Kruskal-Wallis ANOVA, n=20 images in each group. (**D**) Quantitative RT-PCR displays lower expression levels of *Gja1, Scn5a and Kcnj2*, but higher *Hcn4* levels in TBX18-NRVMs with or without A83-01 compared to GFP-NRVM at day 7. n=5 biological replicates in each group, \**P<*0.05, one-way ANOVA with Bonferroni test. (**E**) Fluorescence images: freshly isolated adult rat ventricular myocytes transduced with GFP or TBX18-Zsgreen at 3 weeks post-injection in vivo. No difference was observed in spontaneous beating rates (left bar graph) or the ratio of cell width to length between TBX18-VMs with or without A83-01 treatment. n=70 cells in each group, Kruskal-Wallis ANOVA for beat rate comparison and one-way ANOVA with Bonferroni test for ratio of width/length.

### Inhibition of Tgfβ mitigates loss of syncytial pacing in TBX18-iPMs

To understand the longevity of the *de novo* automaticity, we examined the spontaneous contractions in TBX18-NRVMs in culture. Control GFP-NRVMs monolayers were mostly quiescent, showing occasional contractions that are thought to result from reentrant waves (fig. S10, movie S1) (*23*). In contrast, TBX18-NRVMs monolayers showed rhythmic contractions as previously demonstrated (*6*). However, the rhythmic contractions of TBX18-NRVMs began to waver within the first few days of gene transfer (fig. S10, movie S2). By day 7, TBX18-NRVMs still showed single-cell contractions, but were largely incapable of generating syncytial contractions (fig. S10, movie S2). Measurements of spontaneous field potentials with multi-electrode arrays (MEAs) demonstrated the loss of syncytial automaticity and rhythmic stability in TBX18-NRVMs (Fig. 5A). In contrast, TBX18-NRVMs treated with A83-01 continued to show rhythmic and syncytial contractions and prevented the loss of electrical pacing in TBX18-NRVMs (Fig. 5A & 5B, fig. S11, movie S3 & S4). Measurements of spontaneous intracellular Ca^2+^ transients further confirmed the synchronized excitation-contraction coupling of TBX18-NRVMs beyond week one in the presence of the Tgfβ inhibitor (Fig. 5B, fig. S11C and movie S4).

**Fig. 5.**
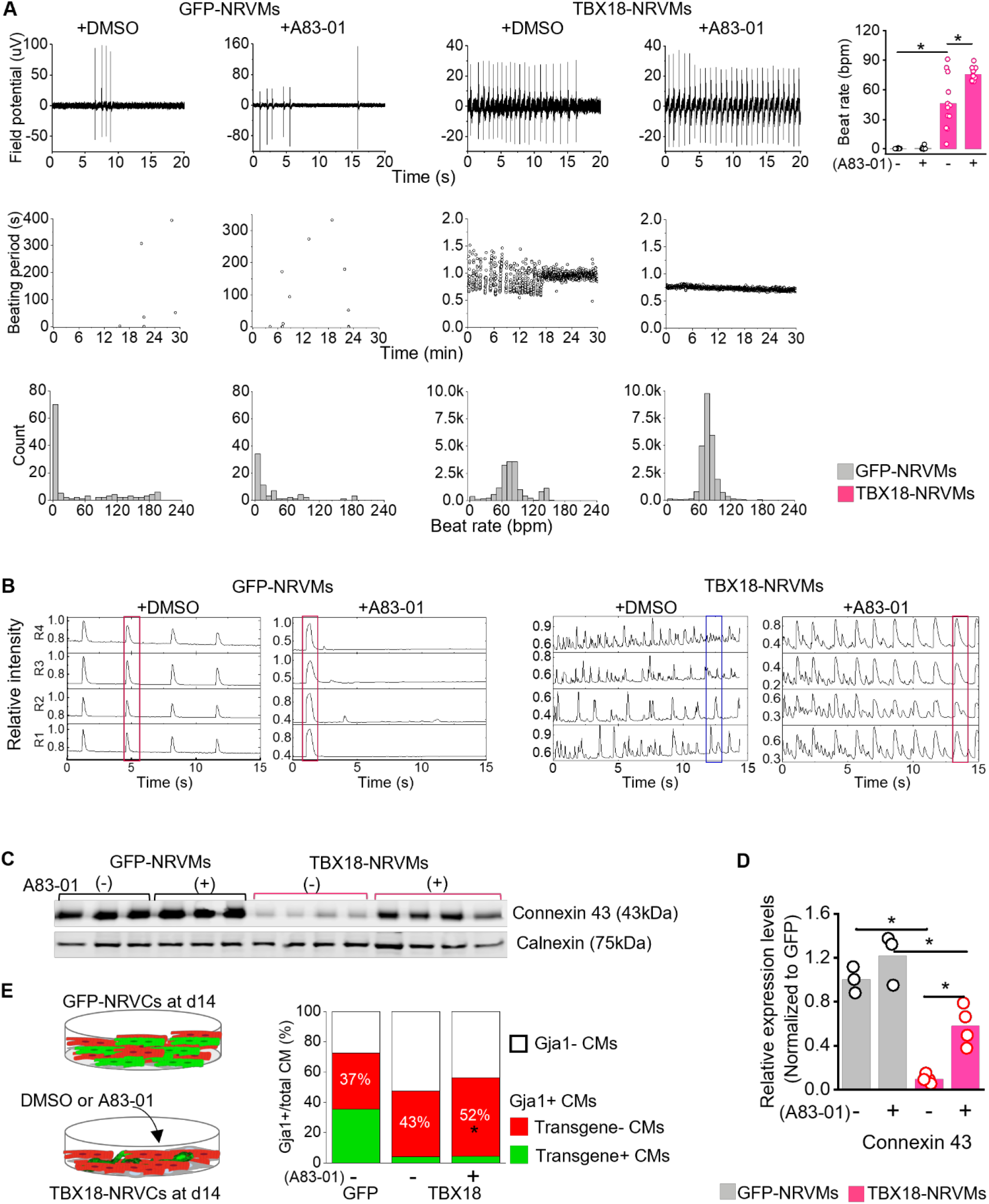
Loss of pacing by TBX18-iPMs is mitigated by Tgfβ inhibition. (**A**) MEA data point illustrating spontaneous electrical activity (top panel), beating period variation (middle panel) and histogram analysis of spontaneous beat rate of the NRVM monolayers at d7. TBX18-NRVMs with A83-01 present faster spontaneous beat rate and more stable beating period. n=10 wells in each group, \**P*<0.05, one-way ANOVA with Bonferroni test. (**B**) Regions of interest (R1 to R4) plotted from spontaneous whole-cell Ca^2+^ transient measurements of NRVMs at d9 illustrates the frequency of spontaneous Ca^2+^ activity and the synchrony of those events. The red bar indicates syncytial beats among multiple regions of interest, but the blue bar indicates asynchronous beats, 28 frames/s. (**C, D)** Cx43 protein levels quantified by western blot analysis were substantially lower in TBX18-NRVMs than in GFP-NRVMs, which is partially restored by A83-01 treatment (n=3 to 4 biological replicates in each group, \**P*<0.05, one-way ANOVA with Bonferroni test). (**E**) The diagrammatic map shows the majority of CMs are TBX18 negative CMs (red color, left) and A83-01 increases the proportion of *Gja1*^+^ CMs in non-transduced CMs by around 10%, with minimal effects on TBX18^+^ CMs (green color, right). \**P*<0.001 compared to TBX18-CMs, Chi-square test.

We carried out similar experiments with NRVCs that contain higher degree of nonmyocytes, including 6.2% vimentin^+^ fibroblasts, 5.7% Pdgfrα^+^ active, and 3.7% Cd31^+^ endothelial cells (fig. S1). Control NRVCs monolayers were spontaneously Tgfβ1 signaling (*24*). Indeed, treatment of control NRVCs monolayers with a Tgfβ inhibitor A83-01 substantially decreased the spontaneous activity at d8(fig. S12A & 12B). In contrast, TBX18-NRVCs monolayers showed faster spontaneous contraction rates compared to control NRVCs, and A83-01 treatment further increased synchronized contractions at d8 (fig. S12A &12B, movie S5). The data indicate that the underlying mechanisms for spontaneous activity and the impact of Tgfβ inhibition are distinct between TBX18-NRVCs and GFP-NRVCs. These results are in line with the flow cytometry data showing that A83-01 treatment lowered the content of vimentin^+^ fibroblasts in TBX18-NRVCs from 25.5±2.9% to 13.9±2.9% at d8, to the level observed in control GFP-NRVCs (n=6 biological samples, *P<*0.05, fig.12C). Together, the data demonstrate that inhibition of Tgfβ signaling with A83-01 can rescue loss of synchronous pacing of TBX18-iPMs by suppressing excessive fibrosis.

Connexin 43 (Cx43)-mediated gap junctions are found in the SAN, but their presence is on the strands of atrial myocytes that intercalate with strands of nodal myocytes and not on the pacemaker cells *per se* (*25*). We have previously demonstrated that re-expression of TBX18 strongly and acutely suppresses Cx43 in ventricular myocytes (*26*). Because the efficiency of direct cell reprogramming is generally very low (1-20%) (*27*) and the proportion of iPMs in TBX18-NRVMs appears to be follow this biology (4.9%, Fig. 4B & 4C), depletion of Cx43 in iPMs alone may not account for the loss of Cx43 expression in TBX18-NRVMs. Tgfβ signaling can act through a paracrine manner and downregulate Cx43 gap junctions (*28*), which suggested that cardiomyocytes that were not transduced with TBX18 may still be effected by Tgfβ signaling. We examined the level of Cx43 expression in the context of Tgfβ inhibition. Inhibition of Tgfβr with A83-01 restored decreased Cx43 protein levels to some extent in TBX18-NRVMs (2.2-folds higher) compared to untreated TBX18-NRVMs (Fig. 5C & 5D, fig. S13). Indeed, restoration of gap-junctional coupling was restricted to cardiomyocytes that were not expressing TBX18, showing inverse correlation between TBX18 expression and Cx43 in TBX18-NRVMs treated with A83-01 (Fig. 5E).

We asked whether cell-cell decoupling and ECM remodeling in TBX18-NRVMs would create a gap between cells and/or compromise attachment to the bottom substrate. Changes in cell attachment and cell-cell coupling can be measured as electrical impedance between two electrodes that are overlaid with cells (*29*). Electrical impedance began to decrease by d3 after TBX18 gene transfer and reached minimal values by d7 while it held steady in GFP-NRVMs for more than one week. Treatment with A83-01 largely mitigated the drop in electrical impedance in TBX18-NRVMs for two weeks (fig. S14), suggesting that the cell-cell physical contact of TBX18-NRVMs and their attachment to the bottom substrate remained steady upon inhibition of Tgfβ signaling. Together, the data indicate that inhibition of Tgfβ1 signaling preserves cell-cell electrical and physical coupling in TBX18-iPMs and surrounding cardiomyocytes. This rescues loss of synchronous automaticity of TBX18-iPMs.

### Transient inhibition of Tgfβ signaling leads to durable ventricular pacing by TBX18 in vivo

Earlier work by us and others indicated that TBX18-induced ventricular pacing began to wane *in vivo* within days after the gene transfer (*7, 8*). We set out to test the hypothesis that the waning of TBX18-induced cardiac pacing can be mitigated by transient inhibition of Tgfβ signaling. To this end, we first generated chronic and severe bradycardia in adult rats by electrically ablating the atrioventricular node one week before transgene delivery (*30*). Upon confirming stable and complete heart block, the animals were randomly assigned into three groups for direct myocardial injection of a transgene (d0, Fig. 6A). Animals in groups 1 and 2 were injected with Adv-TBX18 into the left ventricular anterior apex combined with intraperitonial implantation of an osmotic minipump loaded with either a vehicle only (DMSO, group 1) or A83-01 (2.5 mg/kg/day, group 2) so as to slow-release the drug for the next 7 days. Control animals (group 3) were injected with Adv-GFP and implanted with an osmotic minipump loaded with A83-01 for one week. A GFP-injected group without A83-01 was not included based on our previous data that Adv-GFP alone did not yield ectopic ventricular pacing in animals with heart block (*7, 30*). To quantitatively assess the efficacy and longevity of biological pacing, all animals were implanted with a telemetry for 24/7 real-time recording of electrocardiograms (ECGs) for three weeks after the gene transfer.

**Fig. 6.**
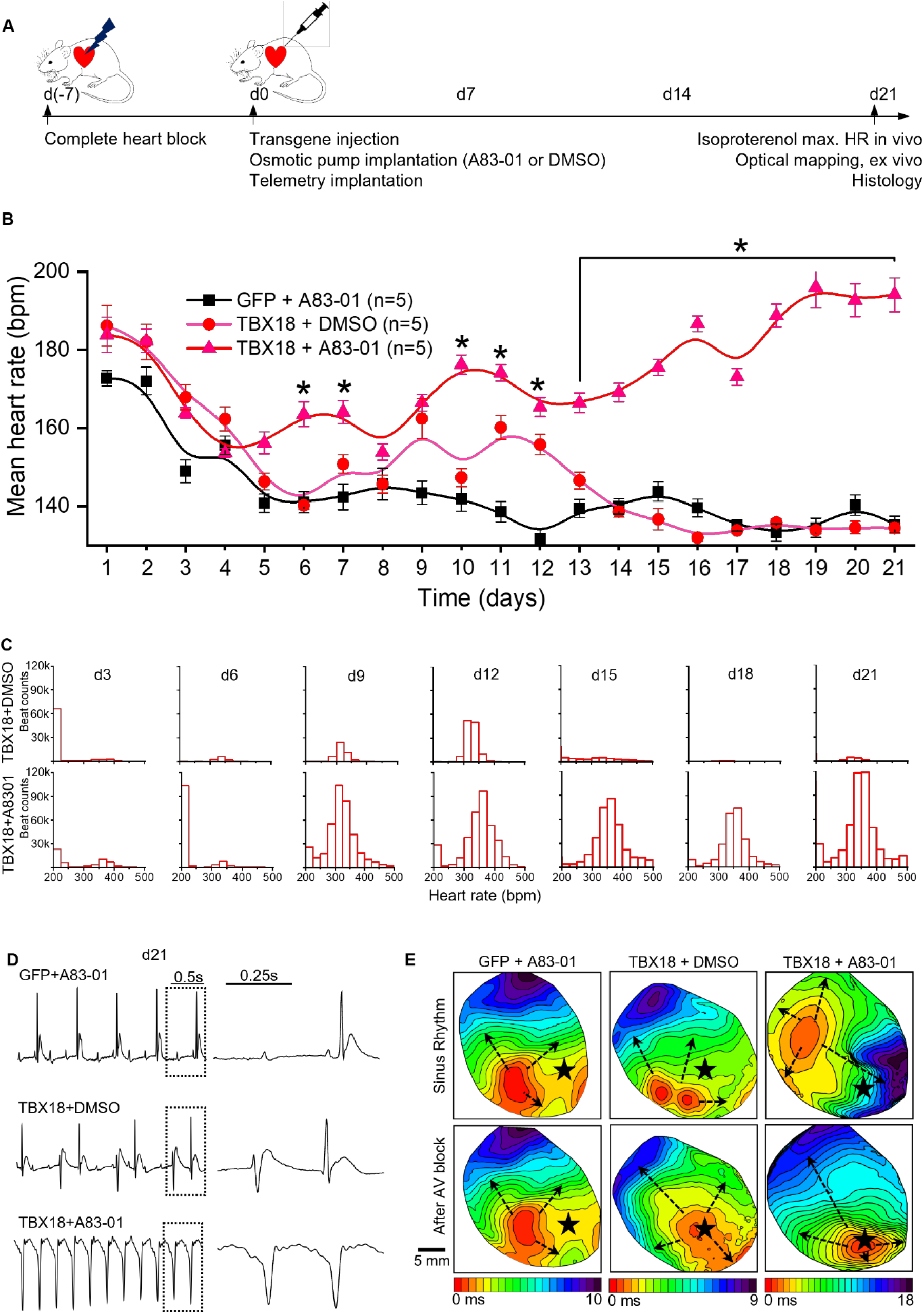
Transient inhibition of Tgfβ rescues the waning of TBX18-induced cardiac pacing in vivo. (**A**) Longitudinal study design for TBX18-induced cardiac pacing in a rat model of CAVB *in vivo* for the data presented in B to D. (**B**) Daily meant heart rates of the animals recorded from 24/7 ECG telemetry for 3 weeks after transgene injection. n=5 animals per group, mean ± SE, \**P<*0.05, one-way ANOVA with Bonferroni test. (**C**) Heart rate histogram of a 24-hour period illustrating the de novo ventricular rate near 350 bpm, which is sustained throughout the 3-week study period in TBX18-injected animals with A83-01. (**D**) Ambulatory ECG traces demonstrate junctional QRS complexes in control animals (upper panel), competition of de novo, wide-QRS complexes with junctional QRS complexes in TBX18-injected animals (middle), and mostly de novo, wide-QRS complexes in TBX18-injected animals with A83-01 (lower panel). (**E**) Optical mapping of membrane potential propagation before (upper panel) and after (lower panel) AV node ablation. A separate group of animals were injected with the transgene (d0), but without AV block. The heart was harvested at d21 after gene delivery, and AV node ablation was performed under retrograde perfusion in normal Tyrode’s solution. After creating the AV block, the earliest electrical activation site shifted to the injection site in the hearts injected with TBX18 plus A83-01 in 5 of 5 animals, and in 3 of 3 TBX18-injected animals (without A83-01) after isoproterenol wash-in. The earliest electrical activation site remained the same in all GFP-injected hearts (5/5) even after isoproterenol wash-in. Star: injection site of transgene injection. Arrows: direction of wave propagation.

During the three-week study period, the GFP control animals showed persistently slow mean HRs, averaging 142.4±3.3 beats per minute (bpm) one week after the gene injection and until the end of the 3-week study period (n=5, Fig. 6B). The mean HRs of the animals injected with Adv-TBX18 without Tgfβ inhibition (Fig. 6B, Adv-TBX18+DMSO) were significantly higher than that of the control animals but began to decline during week 2 and became indistinguishable from the HRs of control animals. In contrast, the mean HRs of the TBX18-injected animals with A83-01 were significantly higher than the HRs of either the control or TBX18-injected animals without A83-01 by d7, and maintained the higher mean HRs for the entire 3-week study period and additional 2-week follow-up (Fig. 6B, fig. S15).

Histogram analysis of daily HRs revealed two major HRs in TBX18-injected animals, a slow HR that is similar to the control animals’ junctional rhythm (121±12 bpm at d10) and a faster HR at 326±26 bpm, presumably induced by TBX18 (Fig. 6C, fig. S16). Over the 3-week study period, the faster HR diminished in TBX18-injected animals. In contrast, animals injected with Adv-TBX18 combined with Tgfβ inhibition sustained the biological pacemaker function with a mean HR of 194±4.3 bpm at d21 (Fig. 6C, fig. S16). We examined HR response to β-adrenergic stimulation with isoproterenol under anesthesia. Animals injected with TBX18 combined with Tgfβ inhibition showed highest increase in HR compared to the other two groups of animals upon isoproterenol injection (fig. S17), indicating highest chronotropic competence. Taken together, the data demonstrate that inhibition of Tgfβ pathway enables durable ventricular pacing by TBX18 *in vivo*.

Examining lead II of the surface ECG of the animals indicated that the QRS complexes of control animals were slow and narrow, typical of a junctional rhythm (Fig. 6D, upper panel). In contrast, TBX18-injected animals showed two types of QRS complexes; slow and narrow ones that are indicative of junctional rhythm and wide and faster QRS complexes that are indicative of ectopic ventricular rhythm (Fig. 6D, middle panel). The wider QRS complexes in both TBX18 with DMSO and TBX18 with Tgfβ inhibitor animals were negative in polarity and in line with the apex injection, designed to generate apex-to-base retrograde conduction. To better understand whether the ventricular pacing in TBX18-injected animals originated from the gene injection site, we examined activation and propagation of the electrical waves in the whole heart *ex vivo* by optical mapping. In this separate experiment, normal adult rats were injected with the transgene at the left ventricular wall, and at d21 the heart was harvested and retrograde-perfused (fig. S18). The earliest activation site remained the same in the GFP-injected control hearts under sinus rhythm and upon AV node ablation (Fig. 6D, left, movies S6). In contrast, the earliest electrical activation shifted to the site of TBX18 injection from remote regions of the ventricles in TBX18-injected heart regardless of the presence/absence of Tgfβ inhibition (Fig. 7D, middle and right, movies S7 and S8). These data indicate that the faster HRs in TBX18-injected animals originated from the gene injection site, in line with their function as a biological pacemaker.

**Fig. 7.**
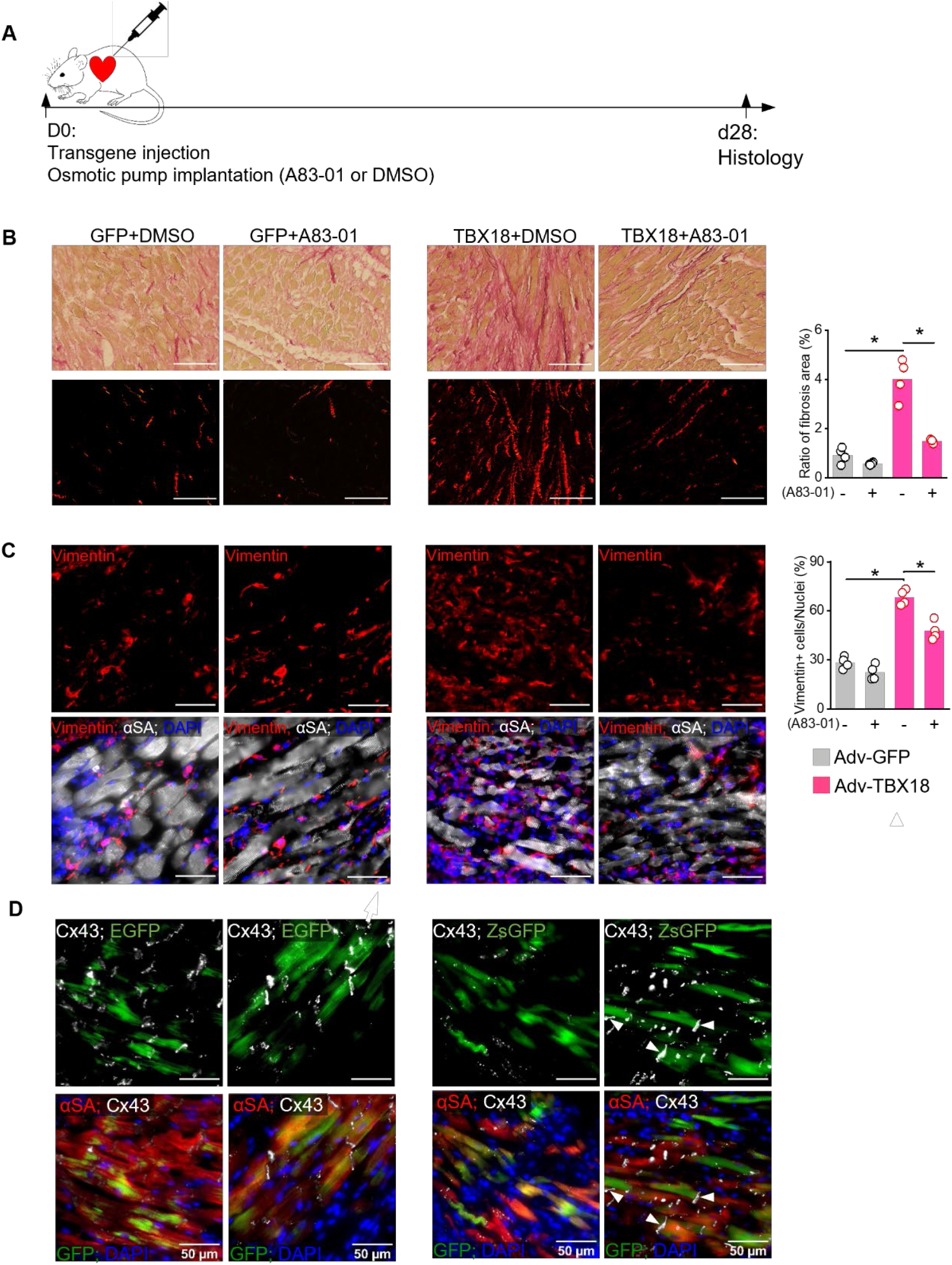
Transient inhibition of Tgfβ signaling ameliorates fibrosis at the site of TBX18 delivery. (A) The schematic design for TBX18-induced histological changes. (**B**) Picrosirius red staining at 4 weeks after transgene injection, treated with A83-01 or DMSO for the first 7 days. This experiment was performed in a separate group of animals without CAVB surgery. Collagen deposits in TBX18-injected animals were substantially reduced with A83-01 treatment (lower panel). n=4 animals per group, \**P<*0.05, one-way ANOVA with Bonferroni test. Scale bar=100 µm. (**C**) Immunostaining of vimentin^+^ (upper panel) at d28 at the transgene injection site. Scale bar=50 µm, n=4 animals per group, \**P<*0.05, one-way ANOVA with Bonferroni test. (**D**) Immunostaining of Cx43 at the injection site at d28. Cx43 expression is barely detectable at the site of TBX18-injection but is restored between TBX18-expressing iPMs and non-transduced myocytes (white arrows) in animals treated with A83-01.

We examined the degree of fibrosis at the transgene injection site. The site of gene delivery was confirmed by the GFP reporter in the left ventricular free wall (Fig. 7A). At d21, TBX18-injected animals treated with A83-01 showed substantial reduction in collagen deposits compared to those without Tgfβ inhibition as revealed by Picrosirius staining (Fig. 7B, fig. S19). Treatment with A83-01 substantially abrogated the increase in vimentin^+^ nonmyocytes found at the site of TBX18 gene injection (Fig. 7C).

Tgfβ-mediated fibrosis in the SAN could lead to sinus node dysfunction (*31*). We examined histological sections at the injection site to understand whether the reduced fibrosis with Tgfβ inhibition might have improved cell-cell electrical coupling *in vivo*. Weak cell-cell electrical coupling insulates the core of the SAN from the hyperpolarizing electrical sink generated in the general myocardium (*25*). However, complete loss of cell-cell electrical coupling would result in failure to capture the neighboring myocardium. Indeed, Cx43 expression were nearly absent at the site of TBX18 injection (Fig. 7D second from right). In contrast, Cx43-mediated gap junctional coupling at the interface between TBX18-iPMs and ventricular myocytes were restored in TBX18-injected animals with Tgfβ inhibition (Fig. 7D right panel). Taken together, the data indicate that TBX18-induced biological pacing elicits *in situ* fibrosis and cell-cell decoupling, leading to loss of pacing. Transient inhibition of Tgfβ signaling suffices to mitigate this loss and achieves durable cardiac pacing.

## DISCUSSION

The SAN is naturally fibrotic (*14, 32*). Similarly, our data indicate that re-expression of exogenous TBX18 led to fibroblast activation and interstitial fibrosis (Fig.1). Unlike the SAN, however, TBX18-triggered fibrosis was overt and overwhelming, leaving the gene transfer site dominated by nonmyocytes and collagen deposits where few cardiomyocytes could be sighted (Fig. 1). The fibrosis appears to be mediated, at least in part, by Tgfβ signaling and phosphorylation of Smad2/3 (fig. S8B). Small molecule-mediated inhibition of Tgfβ signaling softened the fibrosis by downregulating phosphorylation of Smad2/3 (fig. S8B). Functionally, inhibition of Tgfβ signaling sufficed to rescue gradual loss of TBX18-induced automaticity *in vitro* and created durable ventricular pacing *in vivo* for three weeks in a rat model of chronic and complete heart block.

Sinus rhythm is a two-step process of pace-and-drive; the inherent automaticity of the pacemaker cells generates oscillatory electrical stimulation (i.e., pace), which needs to propagate into and depolarize the surrounding myocardium (i.e. drive) (*33*). The unusually high nonmyocyte content within the SAN and the resultant weak electrical coupling protects the SAN from the hyperpolarizing electrical sink of the myocardium. However, too much fibrosis disengages the SAN from the atrial myocardium leading to an exit block (*31, 34*). Our data suggest that biological pacing created by TBX18 gene transfer may succumb to exit block without Tgfβ inhibition (*31, 34*).

A minimal degree of cell-cell electrical coupling would be required for the pacemaker cells to synchronize even in the SAN where the cell-cell electrical coupling is weak (*35*). Gap junctional coupling was significantly downregulated in non-transduced cardiomyocytes as well as among TBX18-transduced cardiomyocytes (Fig. 5E), suggesting that both cell-autonomous and non-cell autonomous mechanisms conspire to significantly downregulate the gap junctions. This notion is supported by the co-culture experiments in which the shared media between TBX18-NRVMs and naïve FBs in a transwell system sufficed to trigger fibrotic signals in the FBs (fig. S8C). This suggests that the fibrosis elicited by TBX18 gene transfer led to proliferation and activation of the FBs in a paracrine manner. The present study does not delineate the mechanism by which the ‘bystander’ cardiomyocytes lose gap junctional coupling due to TBX18-transdueced cardiomyocytes. Nonetheless, the severe downregulation of gap junctions would hinder the small population of iPMs to pace and/or drive. Inhibition of Tgfβ signaling can mitigate the loss of gap junctional coupling in the diseased myocardium (*36, 37*). Here, Tgfβ inhibition with A83-01 partially restored Cx43 expression levels *in vitro* (Fig. 5C-5E and fig. S13), and at the apical-to-apical interface between TBX18-iPMs and neighboring ventricular myocytes *in vivo* (Fig. 7D). This likely have contributed to successful impulse propagation from TBX18-iPMs to the surrounding myocardium.

Our scRNA-seq data indicate that activation of FBs to *Acta2*^+^ myoFBs occurs by d3 after TBX18 gene transfer (Fig. 2). The level of Acta2 expression in myoFBs remained elevated beyond this acute phase (fig. S4), indicating that inhibition of Tgfβ signaling needs to be applied early when TBX18 is actively expressed. The half-life of Tbx18 proteins is brief at 2.5 hours (*38*), suggesting that TBX18 may not linger long after the initial gene transfer. Our *in vivo* proof-of-concept experiments were designed to test the hypothesis that one-week, transient inhibition of Tgfβ signaling suffices to reduce fibrosis and enable long-term biological pacing by TBX18 in a rat model of complete heart block (Fig. 6B & 6C, Fig. 7A-7C). The data support our approach that inhibition of Tgfβ signaling does not need to be extended beyond one week after TBX18 gene transfer in order to reduce the fibrosis long-term.

Estimates for FB content in the adult mouse heart range from 10-20% (*39, 40*). The FB contents in our neonatal ventricles were 5.7% before preplating (NRVCs) and 2.1% after preplating (NRVMs, fig. S1). This may suggest lower nonmyocyte content in the neonatal heart, and/or be due to cell dissociation that was optimized to recover viable myocytes. The scRNA-seq data indicate that a significant population of FBs underwent spontaneous activation to *Acta2*^+^ myoFBs during culture (Fig. 2F). This is in line with previous reports (*24, 41*) and is the likely reason for the elevated level of myoFB content in control conditions *in vitro* (Fig. 2F and 3A). This tendency for spontaneous activation of FBs may have contributed to the higher myoFB content and upregulated fibrosis-related signaling pathways in TBX18-transduced cultures.

Previous studies of TBX18-induced biological pacing showed waning efficacy around one week after the gene transfer which coincides with the rise of intense fibrosis at the TBX18 injection site (Figs. 1A and B). The TBX18-mediated fibrosis was substantially greater than that observed in control animals where inflammatory/immune responses to tissue injury during gene injection and the viral vectors were likely. Hence, it is plausible that treatment with A83-01, a wide-spectrum Tgfβ inhibitor (*17*), had ameliorated fibrotic signals of most causes. All animals showed elevated HR for 3-4 days following the second thoracotomy and gene delivery, potentially as a consequence of the major surgery and systemic elevation of catecholamine levels (*42*).

Somatic cell reprogramming is inherently inefficient (*27*). It is unclear whether incompletely reprogrammed cardiomyocytes may revert back to their original state. The degree and fate of incompletely reprogrammed cardiomyocytes do not appear to have impacted the durability of TBX18-induced biological pacemaker function. Nonetheless, incomplete reprogramming may become relevant to safety aspects such as neoplasm or arrhythmias. This warrants a dedicated investigation.

In conclusion, our study identified overt fibrosis as the reason for the transience of TBX18-induced cardiac pacing and demonstrated that brief inhibition of Tgfβ signaling is necessary and sufficient for attaining a durable biological pacemaker. The strategy overcomes the major impediment to biological pacing and motivates long-term preclinical studies in a large animal model.

## MATERIALS AND METHODS

### Isolation of neonatal rat ventricular myocytes

NRVMs and nonmyocytes were isolated from 1 to 3-day old neonatal Sprague-Dawley rats as previously described (*6*). Briefly. the lower two thirds of the ventricles were harvested to avoid carryover of the atrioventricular nodal cells. Single cell suspensions obtained from trypsin/collagenase II-digested ventricular tissues were subjected to preplating steps to separate neonatal rat ventricular myocytes (NRVMs) from nonmyocytes. Single cell suspensions before the preplating steps were designated as neonatal rat ventricular cells (NRVC). For all *in vitro* experiments, cells were plated at a density of 210,000 cells per cm^2^ of surface area. Replication-deficienty, recombinant adenoviral (Adv) gene transfer *in vitro* was achieved at a multiplicity of infection (moi) of two. The Adv vectors expressed either human TBX18 gene with a green fluorescent reporter (Adv-TBX18-IRES-zsgreen) or an enhanced green fluorescent protein (Adv-GFP) under a CMV promoter (*6*). Media was refreshed every 2-3 days 2% FBS NRVM medium (*6*). For Boyden chamber experiments, NRVMs were seeded on the fibronectin-coated insert at d0, while the non-transduced FBs from the preplating step were passaged once in serum-free NRVM medium and plated at the bottom of a separate 24-well plate at d0. At d1, the inserts seeded with NRVMs were transferred to the 24-well plate seeded with FBs.

### RNA isolation and RT-qPCR

Total RNA was isolated from cultured cells with TRIzol (Thermo Fisher Scientific, Waltham, MA) following the manufacturer’s protocol. cDNA was generated with a PrimeScript® RT reagent kit (Takara, CITY, STATE), and real-time RT-qPCR was performed with a Rotor-Gene SYBR Green PCR kit (Qiagen Inc., Germantown, MD) in a Rotor-Gene® Q real-time polymerase chain reaction machine (Qiagen Inc., Germantown, MD). The PCR primers were designed with Primer Premier 6 (Premier Biosoft Int, Palo Alto, CA, table S1). Relative gene expressions were determined with the 2^-ΔΔCt^ method by normalizing to the housekeeping gene, ribosomal protein L4 (*Rpl4)* (*43*).

### Bulk RNA-seq

RNA libraries were prepared using standard Illumina TruSeq library kits. A PolyA selection step was performed as part of the library protocol to enrich for messenger RNA. Prepared libraries were then sequenced on an Illumina HiSeq 2000 instrument to generate 50-150 nt single end reads. Raw sequence reads were post-processed to remove Illumina adapters/primer sequences. DEG was carried out using DE analysis App (https://yanli.shinyapps.io/DEApp/) to identify differentially expressed genes.

### Single cell RNA-sequencing and analysis

Attached NRVC monolayers were digested with 0.25% trypsin (Cat# LS003707, Corning, NY) for 5 min at 37 °C, and filtered through a 100-μm cell strainer (Fischer Scientific, Waltham, MA) to generate a single-cell suspension. Single cells were resuspended to a concentration of 700-2,000 cells per microliter. Gel beads-in-emulsion (GEMs) were generated with Chromium X (10X Genomics, Pleasanton, CA) and RNA libraries were prepared using the Chromium Next GEM Single Cell 3’ GEM kit, v3.1 (10X Genomics Inc., Pleasanton, CA) with a targeted cell recovery of 4,000-10,000 cells per reaction. Quality control of cDNA and scRNA-seq libraries were performed using the Bioanalyzer High Sensitivity kit (Agilent Technologies Inc., CA). Libraries were sequenced using Novaseq 6000 (Illumina Inc., CA) with a minimum depth of 50,000–65,000 read pairs per cell. The scRNA-seq data were analyzed with Partek Flow (Partek Inc., USA). FASTQ sequencing files were aligned to the custom rattus norvegicus reference genome (Rnor_6.0) with transgene (hTBX18,EGFP and ZsGFP) using STAR-2.7.8a aligner, and the transcript abundance was estimated using the Partek E/M algorithm based on Ensembl release 104 transcript model with default settings. Original data of scRNA-seq were deposited in the NCBI GEO database (accession GSE189815). For the single cell QA/QC, we applied threshold to filter out low-quality cells based on the total UMI count (<2,000 or >6,300), detected gene count (<1,600 or >60,000), and mitochondrial UMI proportion (>40%) (*44*). Read counts per gene in all samples were normalized by log2 counts per million (CPM) and adding 1 for values of zero. We applied top 20 principal components (PCs) to generate t-Distributed Stochastic Neighbor Embedding (tSNE) with a random generator seed of 10 and a perplexity of 30. Differentially expressed genes (DEGs) were analyzed using Partek’s gene specific analysis method, by excluding genes with <10 reads and with an FDR cutoff adjusted *P<* 0.05 (Poisson regression) and fold change > |2|. Gene Ontology (GO) and Bioplanet pathway analysis were performed using Enrichr (https://maayanlab.cloud/Enrichr/) (*45*) with the threshold of overlap≥3, P-value≤0.05. Significantly upregulated or down-regulated genes were used as input data for enrichment analysis.

### Enzyme-linked immunosorbent assay (ELISA) and Western

The concentration of secreted Tgfβ1 ligand levels was measured by taking 3-day conditioned media at d5 of culture with Tgf beta-1/LAP Rat Uncoated ELISA Kit with Plates by following the manufacturer’s protocol (ThermoFisher Scientific, Waltham, MA). The cells were cultured in a 2% FBS media for ELISA. For Western blots, cell lysates were prepared in RIPA lysis buffer (ThermoFisher Scientific, Waltham, MA), and the total protein concentration was measured with a BCA protein assy (ThermoFisher Scientific, Waltham, MA) on a SpectraMax M5 microplate reader (Molecular Devices, Sunnyvale, CA). For SDS-PAGE, 10 µg of protein per sample were mixed with Novex reducing buffer (ThermoFisher Scientific, Waltham, MA) and LDS sample buffer (ThermoFisher Scientific, Waltham, MA). After denaturation, samples were loaded onto a Novex 4-12% Tris-Glycine Mini Gel™ (ThermoFisher Scientific, Waltham, MA). Gels were transferred in a ice/water container at 100 mA onto a PVDF membrane. Nonspecific binding was blocked by incubating the membrane in Intercept™ TBS Blocking Buffer (LI-COR Biosciences, Inc., Lincoln, NE) for one hour. Membranes were probed with primary antibodies overnight a 4°C, followed by incubation with IR-labeled secondary antibodies at room tempertature for 1 hour. Antibodies and blotting conditions used are detailed in table S2. The immunoreactive bands were detected with an infrared imaging system Odyssy CLx and Image Studio™ Software (LI-COR Biosciences, Inc., Lincoln, NE). Band intensities were quantified with ImageJ (https://imagej.nih.gov/ij/).

### Picrosirius red staining and immunofluorescence staining

Rat hearts were harvested and fixed in 10% formalin, embedded in paraffin, and sectioned at 5 μm thickness for picrosirius red staining, which was performed according to the manufacturer’s instructions (Abcam, Waltham, MA). For immunohistochemistry, freshly harvested heart samples were fixed with 4% paraformaldehyde for 12-16 hours, embedded and sectioned at 8 μm thickness. For immunocytochemistry, NRVMs were rinsed with PBS, fixed with 4% paraformaldehyde in PBS for 15 minutes at 4 °C. Antibodies and their concentrations are listed in table S2. Images were acquired with an ORCA-Flash 4.0 sCMOS camera (Hamamatsu Corp, Bridgewater, NJ) connected to a Leica DMi8 wide-field fluorescence microscope (Leica Microsystems, Inc., Buffalo Grove, IL). Image analysis was performed with ImageJ.

### Flow cytometry

NRVCs or NRVMs were dissociated with 0.25% trypsin (Worthington Biochemical Corp., Lakewood, NJ) to yield a suspension of single cells, and were fixed with 4% paraformaldehyde at 4°C for 20 min, permeabilized in 0.1% saponin and 1% BSA in PBS at room temperature for 20 min and incubated with primary antibodies at 4°C for 1 hour. Cells were washed twice with 3% BSA in PBS and incubated with a secondary antibody for 1 hour. Cells were sorted on the FACS Aria II flowcytometer (BD Biocsiences, San Jose, USA). Data were analyzed with FlowJo software (version 7.6.5; Tree Star, Inc., Ashland, OR).

### Live cell recordings of contraction rates and intracellular Ca^2+^ transients

Plates seeded with NRVMs were incubated in a stagetop incubator at 37°C with 5% CO2 (TokaiHit, Shizuoka, Japan) on a DMi8 wide-field fluorescence microscope (Leica Microsystems, Inc., Buffalo Grove, IL). Spontaneous contractions were recorded real-time under phase contrast microscopy, and synchronous vs. asynchronous contractions of NRVMs were analyzed with a commercial cardiomyocyte contractility analysis software, Pulse (https://pulsevideoanalysis.com) (*46*). For fluorescence recording of spontaneous intracellular Ca^2+^ cycling, cells were incubated with 5 μmol/L Cal-590™ AM (AAT Bioquest, Sunnyvale, CA) with 0.02% Pluronic F127 for 90 min at 37°C, and then were incubated in phenol-free 2% FBS NRVM medium at room temperature for 30 minutes. Real time, time-series image acquisition was performed with an ORCA-Flash 4.0 sCMOS camera (Hamamatsu Corp, Bridgewater, NJ) connected to a Leica DMi8 wide-field fluorescence microscope (Leica Microsystems, Inc., Buffalo Grove, IL).

### Multi-electrode array recording and analysis

MEA plate was coated with fibronectin (50 μg/mL) for 1 h at 37 °C. NRVMs were seeded on standard 96-well MEA plates (Multi Channel Systems, Reutlingen, Germany) with 3 electrodes per well. Extracellular field potential recording *in vitro* was performed using a Multiwell-MEA-System (Multi Channel Systems, Reutlingen, Germany) at 37 °C and 5% CO2 for 30 minutes. Output signals were digitized at 5 kHz. Data were acquired using Multiwell-Screen and analyzed off-line with Multiwell-Analyzer (Multi Channel Systems, Reutlingen, Germany). Beating rates were calculated by taking the total number of synchronized beats normalized to the recording duration.

### In vivo gene transfer in a rat model of complete atrioventricular block

Adult Sprague-Dawley rats weighing 250 to 300 g (Charles River Laboratories, Wilmington, MA) underwent right lateral thoracotomy followed by atrioventricular node (AVN) ablation as previously described (*30*). The animals were followed for one week (d7) with surface ECG to confirm chronic and complete atrioventricular block (CAVB). Upon validating a chronic CAVB, the animals were randomly assigned into three groups and underwent a partial left thoracotomy in the fourth intercostal space for somatic gene transfer. Adv-TBX18-IRES-ZsGreen or Adeno-GFP was directly injected into the left ventricular anterior myocardium at a titer of 1 × 10^8^ fluorescence forming units (ffu) in 100 µl of virus storage buffer composed of 1M sucrose, 10mM KPBS and 0.5% β-cyclodextrin (Sigma-Aldrich, St. Louise, MO), at pH 7.4 (*26, 30, 47–49*). A mini-osmotic pump (Alzet 2001, DURECT Corporation, Cupertino, CA) was implanted in the abdominal cavitiy for one-week infusion of either DMSO alone or A83-01 (MedChemExpress LLC, Monmouth Junction, NJ) dissolved in DMSO at a rate of 2.5mg/kg/day(*30*).

### Telemetry measurements and analysis of electrocardiograms

24/7 continuous telemetry recording of ECGs from ambulatory animals was performed for three weeks as described previously (*30*). Briefly, a dual-lead, biopotential telemeter (TR50BB, Kaha Sciences, Dunedin, New Zealand) was implanted. Lead I electrodes were positioned on the left and right latissimus dorsi muscles; while Lead II electrodes were positioned between the suprasternal fossi and left lower thoracic cage. The telemeter body was implanted in the abdominal cavity. The electrical signals were transmitted to SmartPad (TR181, Kaha Sciences, Dunedin, New Zealand) and were analyzed with LabChart Pro (ADInstruments, Colorado Springs, CO).

### Primary adult cardiomyocyte isolation

To examine the effects of Tgfβ inhibition with A83-01 on cell morphology and automaticity of freshly isolated ventricular cardiomyocytes, normal adult Sprague-Dawley rats weighing 250 to 300 g were injected with the Adeno-TBX18-Zsgreen or GFP at the anterior wall of left ventricle and/or were received mini-osmotic pump (Alzet 2001, DURECT Corporation, Cupertino, CA) implantation for infusion of A83-01 for 7 days. All the virus and A83-01 were used the same dose as described above. The heart was harvested at d21 and ventricular myocytes were enzymatically dissociated using a method previously described (6). Isolated single cardiomyocyte was incubated in physiological Tyrode’s solution containing (mM) 130 NaCl, 5.4 KCl, 1.8 CaCl_2_, 1.0 MgCl_2_, 10 D-glucose, and 10 HEPES, pH 7.4 at 37°C, and used within 6–8 h. Morphometric assays were performed with ImageJ by measuring cell area and cell length (long-axis) of each cardiomyocytes, as described previously (*6*). Then, an average cell width was calculated by dividing the cell area by longitudinal cell length. Spontaneous contractions of cardiomyocytes were recorded real-time under phase contrast microscopy using a DMi8 wide-field fluorescence microscope (Leica Microsystems, Inc., Buffalo Grove, IL).

### Ex vivo optical mapping

To understand whether the ventricular pacing in TBX18-injected animals originates from the gene injection site, we performed optical mapping of sarcolemma voltage signals in the whole heart ex vivo. In this separate experiment, normal adult rats were injected with the transgene at the anterior wall of left ventricle, and at d21 the heart was harvested and retrograde-perfused through the aorta in a Langendorff perfusion system (Radnoti Glass Technology, Monrovia, CA, USA) with Tyrode’s solution containing (mM): 140.0 NaCl, 1.0 MgCl_2_, 5.4 KCl, 1.2 NaH_2_PO_4_, 1.0 HEPES, 11 D-glucose and CaCl_2_ (pH 7.4), gassed with O2. Constant flow perfusion was set to 9.8 mL/min with a peristaltic pump. Hearts were placed in a water-heated chamber to maintain temperature at 37 ± 0.2°C and then 5 μmol/L blebbistatin was added to perfusate to reduce movement artefact. Hearts were stained with voltage sensitive indicator di-4-ANNEPS (Thermo Fisher Scientific, Waltham, MA, USA), using 20 μL of stock solution (1 mg/mL of DMSO) delivered through a bubble trap, above the aortic cannula. The ECGs were continuously monitored with a Powerlab system (AD Instruments, Colorado Springs, CO, USA). Fluorescence images of APs were recorded from the anterior surface of the heart using a CMOS camera (256 × 256 pixels, 1818 frames/s, 2.0 × 2.0 cm^2^ field of view; MICAM03; SciMedia, Costa Mesa, CA, USA). The data were analyzed with a custom-built software program developed in Interactive Data Language (Harris Geospatial Solutions, Broomfield, CO, USA). The activation time points at each pixel were determined from fluorescent (F) signal by calculating the local maximum of dF/dt. Fluorescent images were filtered using a spatial bilateral filter (9 × 9 pixel), and first derivatives were calculated using a temporal polynomial filter (3rd order, 13 points).

### Statistics

Data are expressed as mean ± SD unless otherwise indicated. The assumptions of normality and homogeneity of variance were tested by Shapiro-Wilk tests (p>0.1) and Levene’s tests (p>0.1), respectively. Unpaired Student’s t-test for direct comparisons and one-way analysis of variance (ANOVA) with subsequent Bonferroni multiple comparison tests were performed to determine significant differences among individual means. In cases of unequal variance or nonnormal distribution, nonparametric Mann-Whitney two-sample tests or Kruskal-Wallis were used. Statistical significance was defined as *P<*0.05. All statistical analyses were performed using SPSS statistical software version 19.00 (IBM, Armonk, NY) or Origin Pro (OriginLab Corp., Northampton, MA).

## Supporting information

Supplemental figures with legend

## Ethics statement

All animal studies were performed according to the protocols approved by the institutional animal care and usage committee (IACUC) at the Emory University School of Medicine, and in agreement with the guidelines for federal research published by the National Institutes of Health.

## SUPPLEMENTARY MATERIALS

Titles of Supplementary Figures, Tables and Movies

Fig. S1. Flow cytometry analysis of cardiomyocyte and nonmyocyte populations in NRVMs and NRVCs.

Fig. S2. TBX18 increases cardiac fibroblasts (FBs) and extracellular matrix (ECM) gene expression.

Fig. S3. Fibroblast-related gene expression profiles in single-cell populations of cardiomyocytes (CMs) and fibroblasts (FBs) at d3.

Fig. S4. Tgfβ inhibition mitigates myoFBs activation.

Fig. S5. A83-01 mitigates Tgfβ signaling and inflammatory response in TBX18-transduced cardiomyocytes.

Fig. S6. Tgfβ inhibitor A83-01 suppresses interferon signaling in CMs induced by TBX18.

Fig. S7. Expression of Tgfβ ligands and receptors in TBX18-NRVMs.

Fig. S8. Inhibition of Tgfβ signaling attenuates myofibroblast activation.

Fig. S9. The proportion of transgene^+^ CMs and nonmyocytes (non-CMs) at d3 and d14.

Fig. S10. TBX18-NRVMs gradually acquire higher automaticity but lose synchrony.

Fig. S11. TBX18-NRVMs with A83-01 result in more synchronous beats compared to TBX18-NRVMs alone.

Fig. S12. Treatment with A83-01 rescues asynchronous beats of TBX18-NRVC at d8.

Fig. S13. Inhibition of Tgfβ with A83-01 partially restores reduced connexin 43 by TBX18 in non-reprogrammed myocytes after 7 days.

Fig. S14. Tgfβ inhibition mitigates the drop in electrical impedance in TBX18-NRVMs for 2 weeks.

Fig. S15. Animals with Adv-TBX18 plus A83-01 present persistently biological pacing after 5 weeks.

Fig. S16. Temporal change of heart rate distribution among all ambulatory rats.

Fig. 17. The response of TBX18-created biological pacing to isoproterenol challenge test.

Fig. S18. The schematic diagram of adenovirus injection at the left ventricle and the expression of reporter genes for optical mapping.

Fig. S19. Fibrotic response at the TBX18-Zsgreen or GFP positive area at the edge of the injection site.

Table S1. List of primers used in this study for real-time qRT-PCR.

Table S2. List of antibodies used for immunofluorescent staining, immunoblotting, and flow cytometry.

Movie S1. Typical spontaneous beating pattern and temporal changes of GFP-NRVMs.

Movie S2. Typical spontaneous beating pattern and temporal changes of TBX18-NRVMs.

Movie S3. TBX18-NRVMs with A83-01 result in more synchronous beating compared to TBX18-NRVMs alone (bright field video recording, d7).

Movie S4. TBX18-NRVMs with A83-01 display more synchronous calcium transient compared to TBX18-NRVMs alone (Calcium transient recording with Cal-590™ AM, d9).

Movie S5. Treatment with A83-01 rescues asynchronous beats of TBX18-NRVC (Calcium transient recording, d8).

Movie S6. The activation pattern before and after atrioventricular node ablation in GFP-treated heart.

Movie S7. The earliest activation site shifts to the injection site before and after atrioventricular node ablation in TBX18-treated heart.

Movie S8. The earliest activation site shift to the injection site before and after atrioventricular node ablation in TBX18-treated hearts with A83-01.

## Acknowledgments

We thank Sandra Grijalva and Minji Shin for their assistance in technical aspects of experiments. We thank Dr. Lyra Griffith at the Emory Integrated Genomics Core for sample preparation for the scRNA sequencing experiments and Yerkes Genomics Core for RNA sequencing.

## Funding

National Institutes of Health R01HL143065 (HCC)

National Institutes of Health R01HL147270 (HCC)

National Institutes of Health R01HL157363 (HCC)

National Institutes of Health F31HL149272 (DW)

National Institutes of Health F31HL137367 (SIG)

American Heart Association 20TPA35260085 (HCC)

American Heart Association 19POST34460039 (JF)

American Heart Association 19POST34450268 (NKK)

National Science Foundation NSF1609831 (HCC)

National Science Foundation #2016199765 (DW)

National Science Foundation (SIG)

Department of Defense CDMRP GRANT12901705 PR191598 (HCC)

## Author contributions

Conceptualization: HCC, JF, NKK

Methodology: JF, NKK, NF, TYK, JL, DW

Investigation: JF, NKK, NF, TYK

Visualization: JF, NKK, TYK, NF

Funding acquisition: HCC, JF, TYK

Project administration: JF, NKK, HCC

Supervision: HCC

Writing – original draft: JF

Writing – review & editing: JF, HCC

## Data and materials availability

All data associated with this study are present in the paper or the Supplementary Materials. Original data of scRNA-seq were deposited in the NCBI GEO database (accession GSE189815). Primary data are available in data file S1-7.

Downloading link https://1drv.ms/u/s!AgwEX2a0aSEFhCa-KM41Hk3lcKXd?e=zMfHSt

Data S1. QA-QC of single cell count and the identification of main cell population

Data S2. Original western blot bands

Data S3. Gene ontology and pathway analysis

Data S4. Partek GSA test DEG between TBX18-FB vs GFP-FB Day 3

Data S5. Partek GSA test DEG between TBX18-CM vs GFP-CM Day 3

Data S6. Partek GSA test DEG among TBX18-FB vs. GFP-FB vs. TBX18-FB plus A8301 Day 14

Data S7. Partek GSA test DEG among TBX18-CM vs. GFP-CM vs. TBX18-CM plus A8301 Day 14

