## Supplemental figures with legend for "Inhibition of Tgfβ signaling enables durable ventricular pacing by TBX18 gene transfer"

**Supplementary Figures**

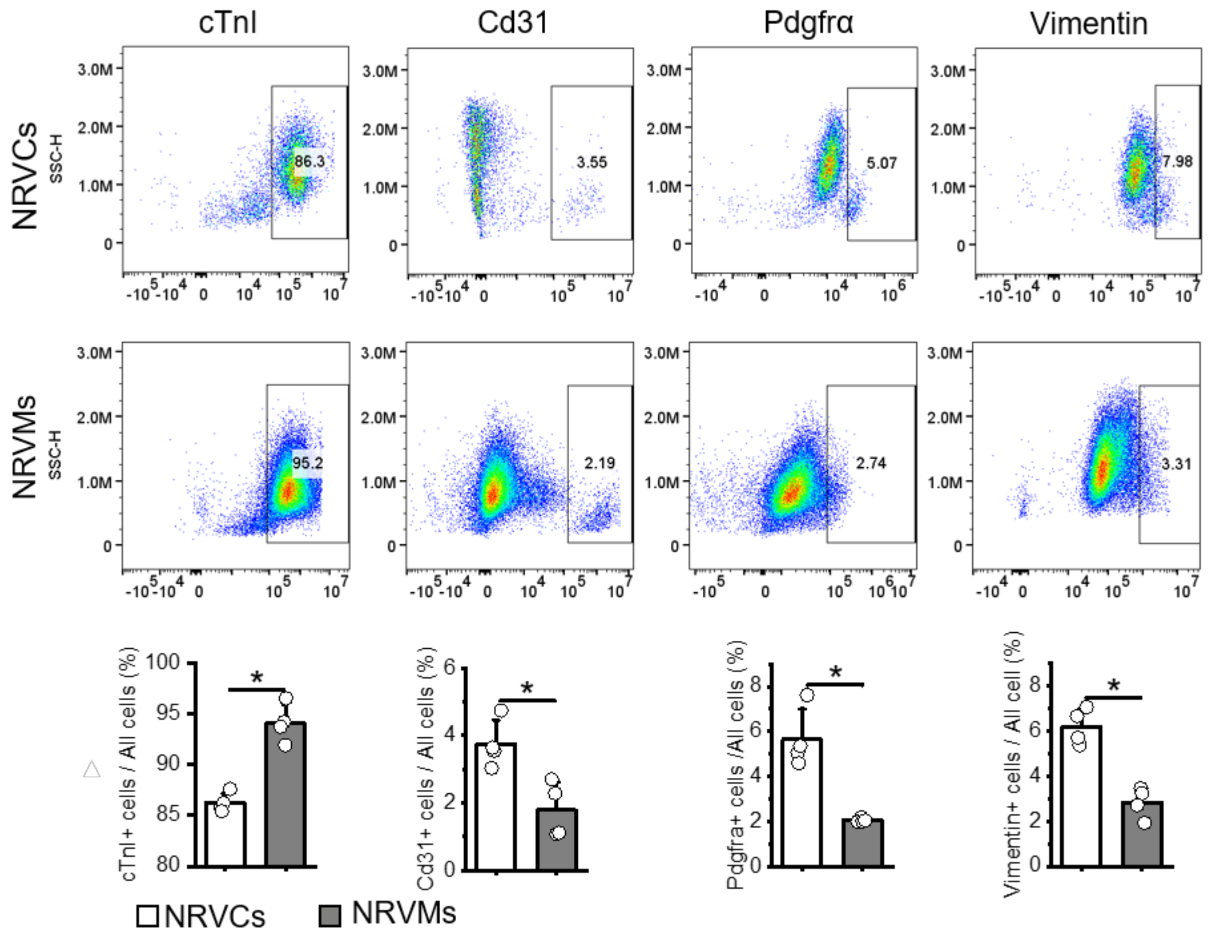


**Fig. S1. Flow cytometry analysis of cardiomyocyte and nonmyocyte populations in NRVMs and NRVCs.** Freshly isolated cells from the neonatal rat ventricles before (NRVCs) or after preplating (NRVMs) were fixed and stained with cell-specific antibodies. cTnI, a cardiomyocyte marker; Cd31, an endothelial cell marker; Pdgfrα, a fibroblast marker; Vimentin, a pan-mesenchymal cell marker. The cardiomyocyte population is higher in NRVMs while the nonmyocyte (non-CM) populations are higher in NRVCs. n=4 biological replicates, Student's t-test.

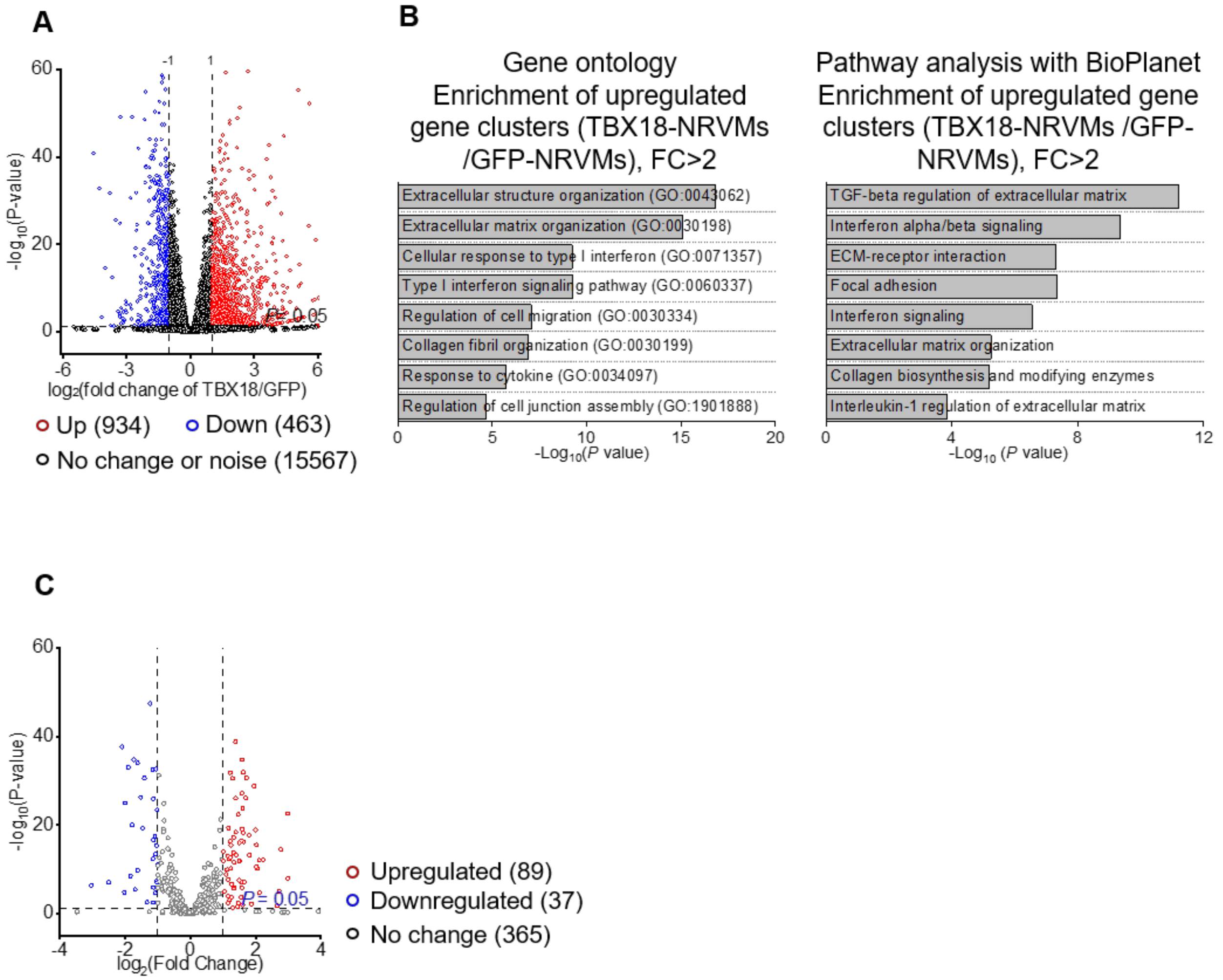


**Fig. S2. TBX18 increases cardiac fibroblasts (FBs) and extracellular matrix (ECM) gene expression.** Bulk RNA-seq data on NRVMs transduced with either Adv-TBX18 or Adv-GFP at d4 after gene transfer. (**A**) Volcano plot of differentially expressed genes (DEG), presented as log2 of fold change in TBX18-NRVMs normalized to GFP-NRVMs. Threshold=2-fold change (FC), and $P<0.05$. (**B**) Gene ontology analysis (left) shows that enriched gene clusters in TBX18-NRVMs are involved in ECM organization and cytokine response. Bioplanet pathway analysis (right) shows that the upregulated gene clusters in TBX18-NRVMs were involved in Tgfβ signaling, interferon signaling, and collagen biosynthesis. (**C**) Volcano plot of DEGs related to Tgfβ regulation of ECM.

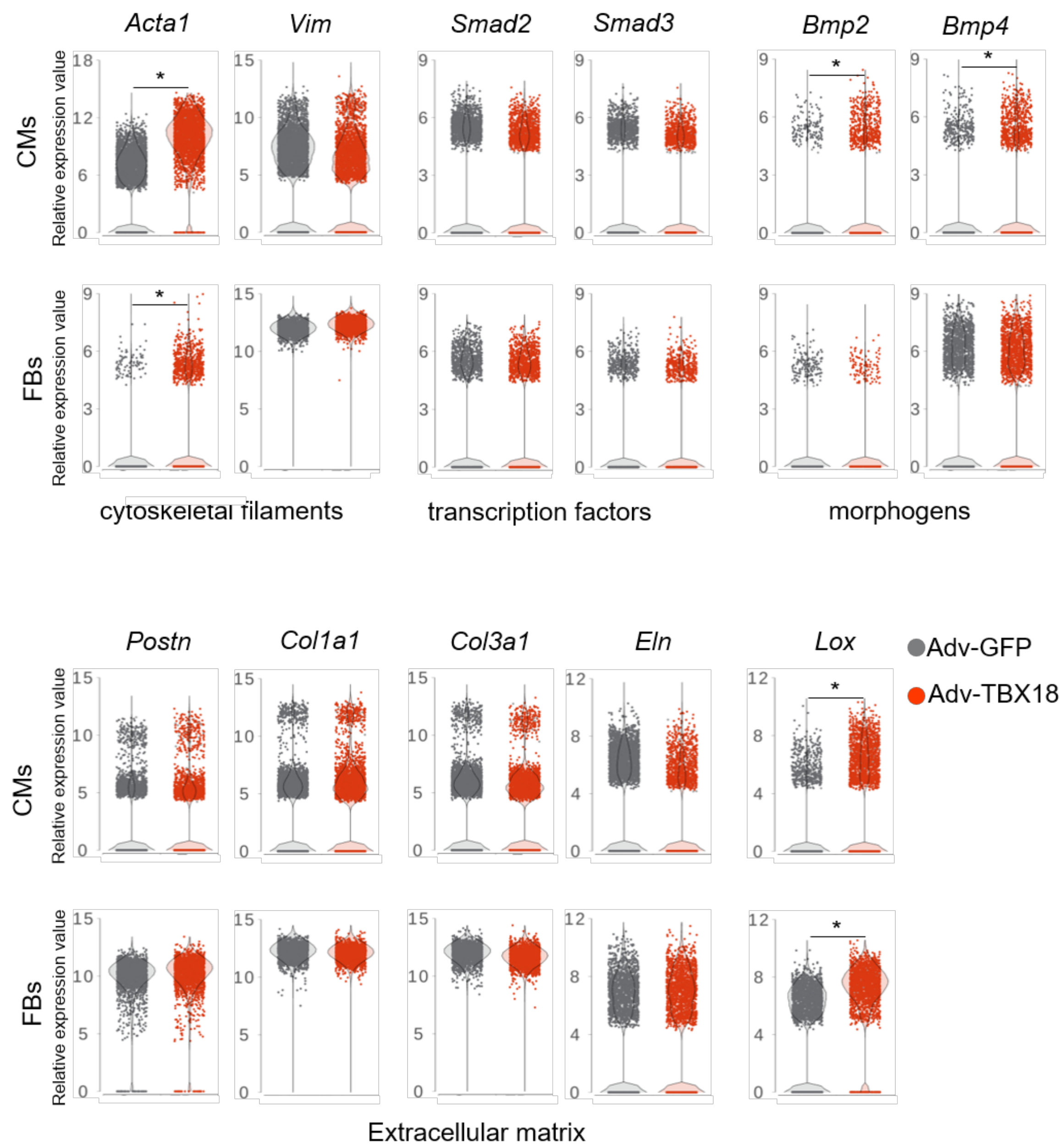


**Fig. S3. Fibroblast-related gene expression profiles in single-cell populations of cardiomyocytes (CMs) and fibroblasts (FBs) at d3.** *$P<0.001$ and $|FC|>1.5$, Partek gene set analysis (GSA) algorithm.

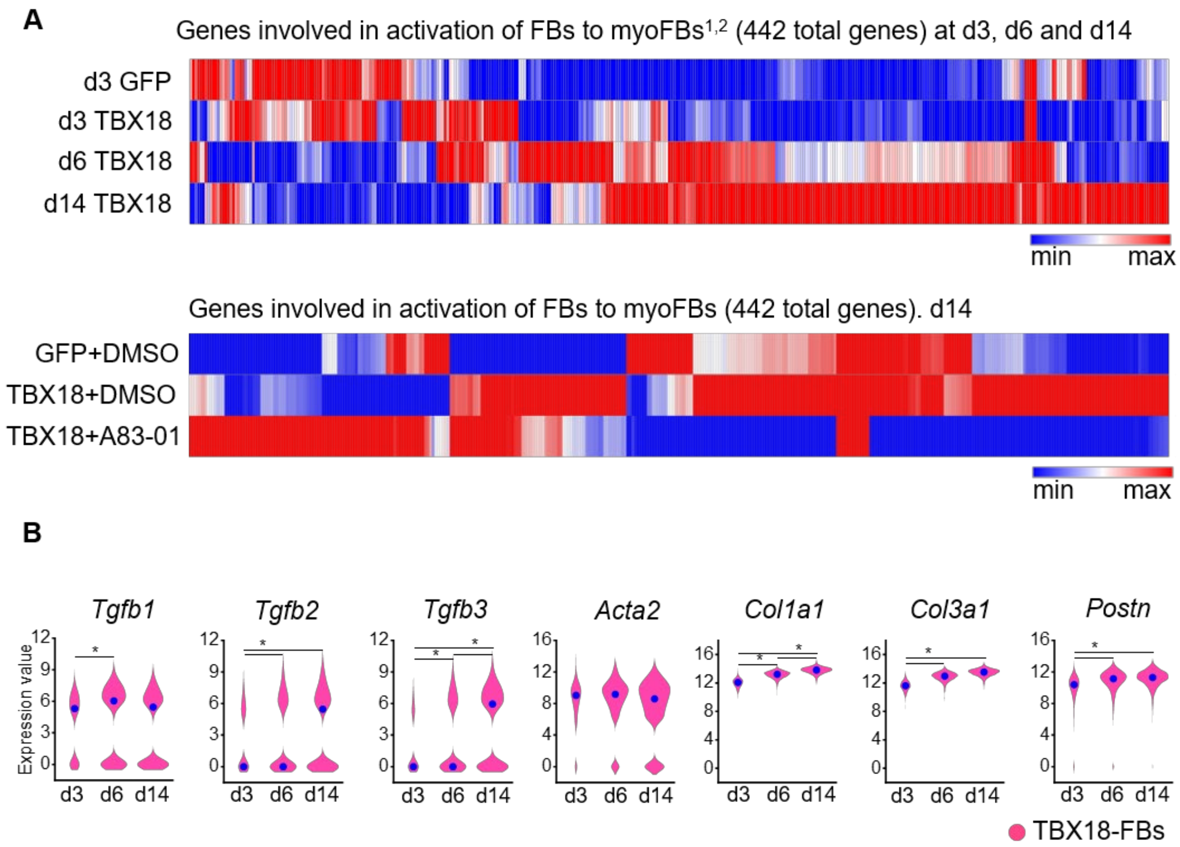


**Fig. S4. Tgfβ inhibition mitigates myoFBs activation.** (**A**) Upper panel: A heatmap of gene expression levels in TBX18-FBs or GFP-FBs with hierarchical clustering of a myoFB gene set. The heatmap highlights the gradual increase in myoFB-related gene expression in TBX18-transduced FBs. Activation of FBs to myoFBs is underway by d3 after TBX18 transduction. Lower panel: A d14 heatmap indicates treatment with A83-01 largely reverses the up and downregulated gene clusters in TBX18-FB. (**B**) Temporal changes of myoFB-related gene expression in TBX18-FBs at days 3, 6, and 14. **P*<0.001 and |FC|>1.5, Partek GSA algorithm.

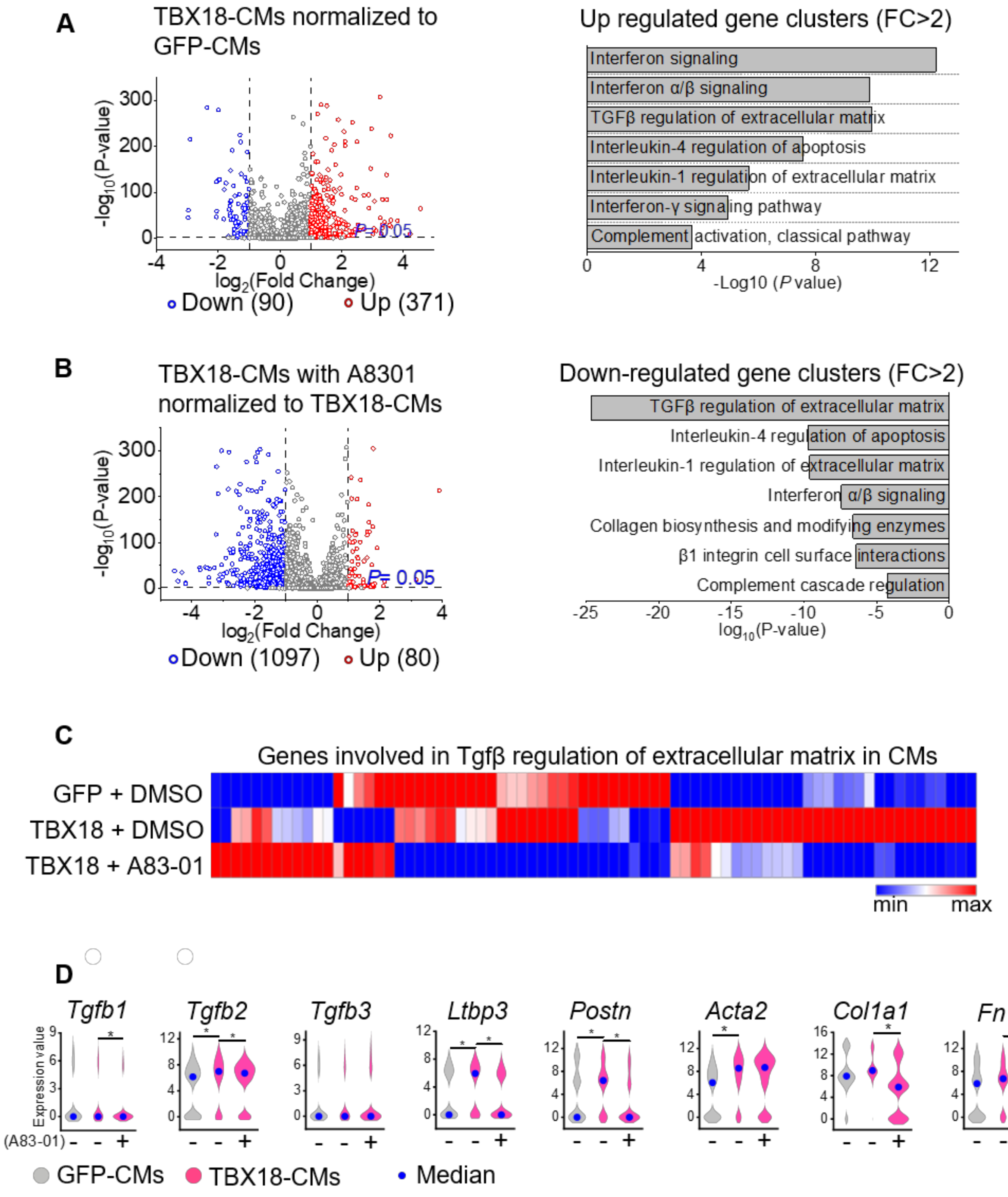


**Fig. S5. A83-01 mitigates Tgfβ signaling and inflammatory response in TBX18-transduced cardiomyocytes.** (**A**) A volcano plot of DEGs in TBX18-CMs normalized to GFP-CMs with a threshold of |FC|>2 and *P*<0.05 (left). Pathway analysis (right) shows upregulated gene clusters in TBX18-CMs, which highlights Tgfβ signaling and cytokine-mediated inflammatory response. (**B**) Volcano plot (left) of DEGs in TBX18-CMs treated with A83-01, normalized to TBX18-CMs with the threshold of |FC|>2 and *P*<0.05. Pathway analysis (right) shows downregulated gene clusters in TBX18-CMs treated with A83-01, which highlights Tgfβ signaling and cytokine-mediated inflammation (right). (**C**) Hierarchical clustering of genes related to Tgfβ regulation of

ECM in CMs demonstrates that Tgfβ signaling is activated by TBX18, which is mitigated by treatment with A83-01. (**D**) Gene expression of Tgfβ ligands and main profibrotic genes are upregulated in TBX18-CMs. Treatment with A83-01 suppresses the increase in Tgfβ-mediated profibrotic gene expression. **P*<0.001 and |FC|>1.5, Partek GSA algorithm.

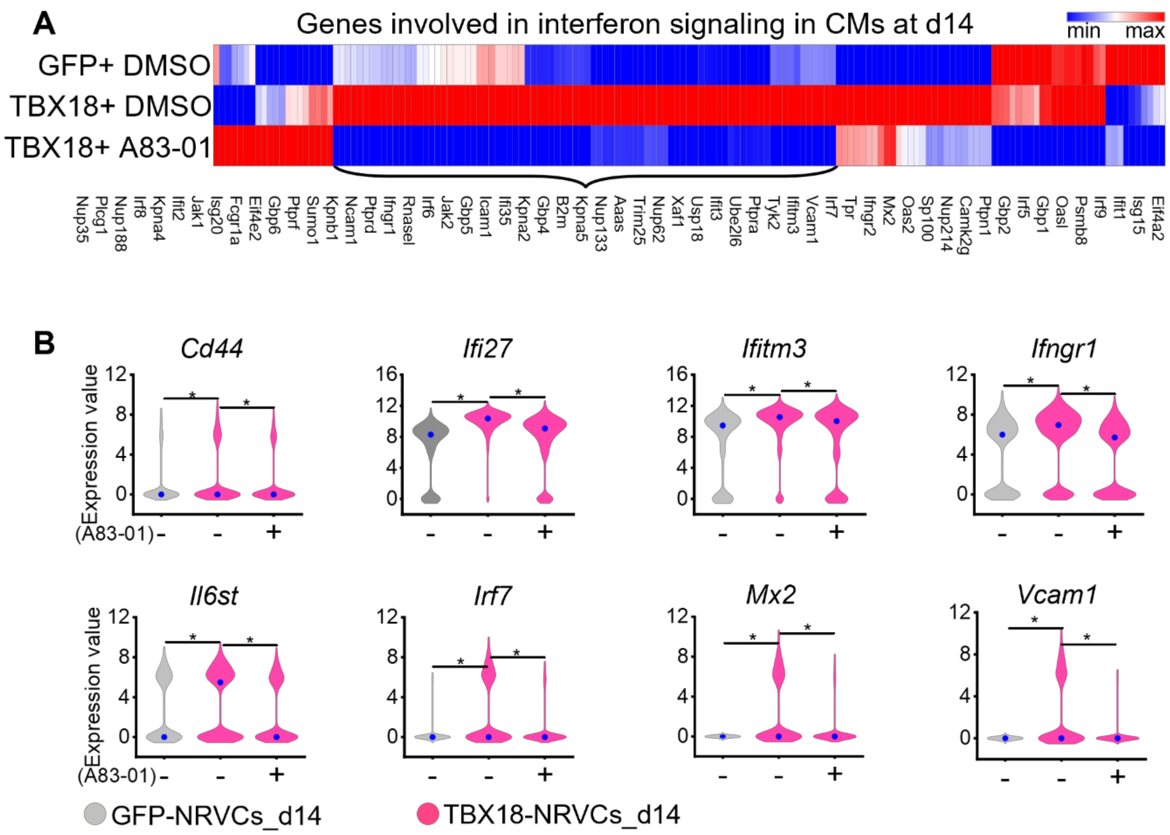


**Fig. S6. Tgfβ inhibitor A83-01 suppresses interferon signaling in CMs induced by TBX18.** (**A**) A heatmap of genes involved in interferon signaling in CMs transduced with either GFP or TBX18. Treatment with A83-01 suppresses a set of genes related to interferon signaling induced by TBX18. (**B**) The expression profile of genes related to interferon signaling and inflammatory response in CMs, *$P<0.001$ and $|FC|>1.5$, Partek GSA algorithm.

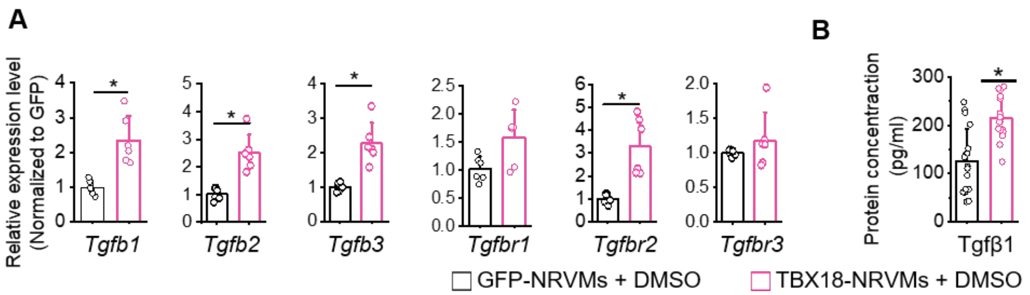


**Fig. S7. Expression of Tgfβ ligands and receptors in TBX18-NRVMs.** (**A**) mRNA transcript levels of Tgfβ ligands and receptors were quantified by RT-qPCR and compared between TBX18- and GFP-NRVMs at d7, n=6 biological replicates for each group. (**B**) Protein levels of the Tgfβ1 ligand were measured in the conditioned medium of TBX18-NRVMs relative to the control medium with ELISA at d5, n=17 replicates in each group. $^{*}P<0.05$, Student's t-test.

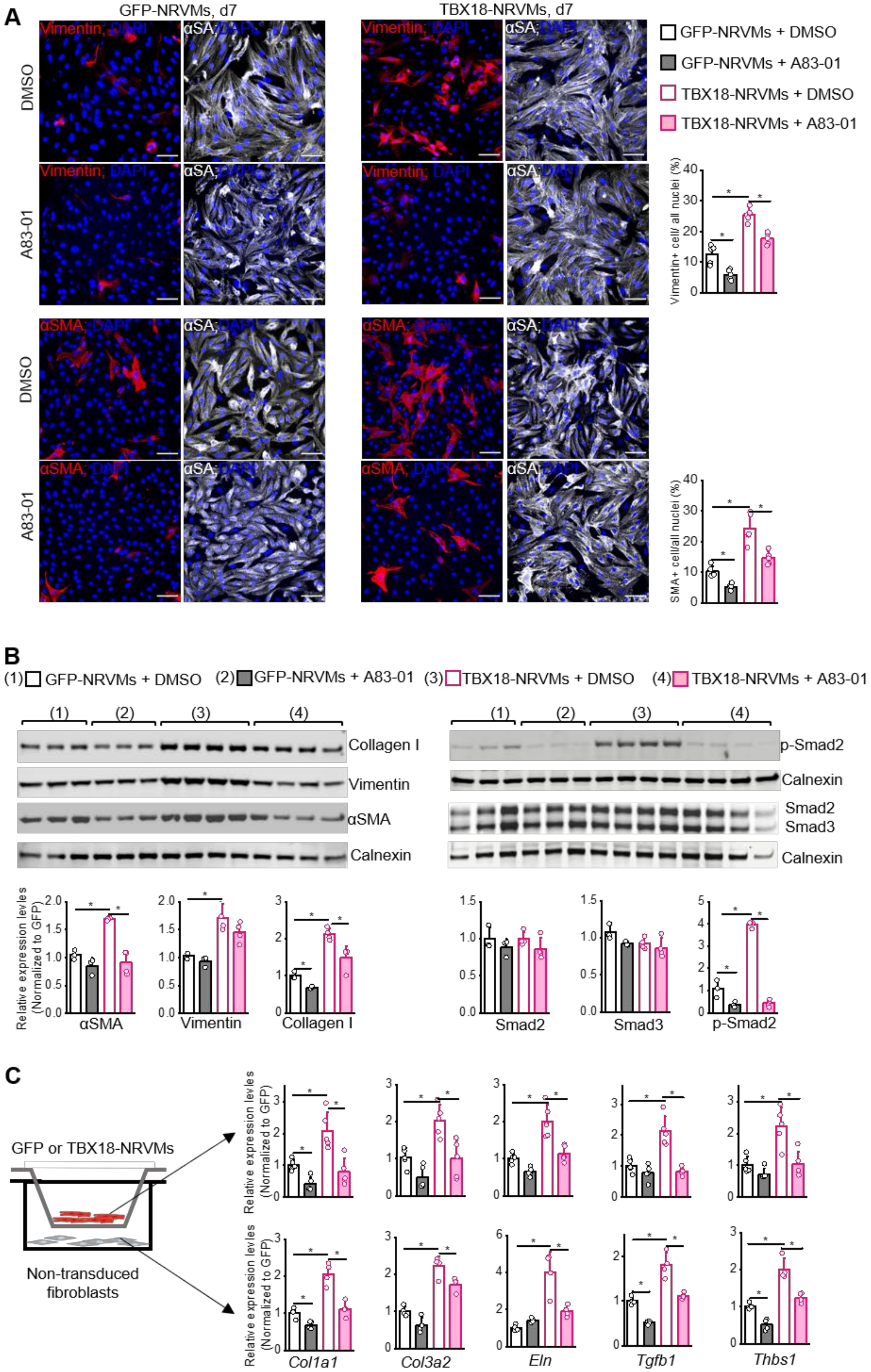
A
GFP-NRVMs, d7
TBX18-NRVMs, d7
DMSO
A83-01
Vimentin; DAPI
αSA; DAPI
αSMA; DAPI
GFP-NRVMs + DMSO
GFP-NRVMs + A83-01
TBX18-NRVMs + DMSO
TBX18-NRVMs + A83-01
Vimentin+ cell/ all nuclei (%)
SMA+ cell/all nuclei (%)
B
(1) GFP-NRVMs + DMSO
(2) GFP-NRVMs + A83-01
(3) TBX18-NRVMs + DMSO
(4) TBX18-NRVMs + A83-01
Collagen I
Vimentin
αSMA
Calnexin
p-Smad2
Smad2
Smad3
Relative expression levels (Normalized to GFP)
C
GFP or TBX18-NRVMs
Non-transduced fibroblasts
Relative expression levles (Normalized to GFP)
Col1a1
Col3a2
Eln
Tgfb1
Thbs1

**Fig. S8. Inhibition of Tgfβ signaling attenuates myoFBs activation.** (**A**). Immunostaining of αSMA$^+$ and vimentin$^+$ cells after 7 days in NRVMs. TBX18-NRVMs exhibit a higher proportion of αSMA$^+$ cells after 7 days *in vitro*; but treatment with A8301 can attenuate the effects of TBX18, n=4 biological replicates, scale bar=50μm. (**B**) Quantitative western blot analysis shows significantly increased expression levels of FB-related proteins in TBX18-NRVMs, for instance, αSMA and collagen I, and phosphorated Smad2 (pSmad2), n=3 to 6 biological replicates in each group, *$P$<0.05. (**C**) Quantitative RT-PCR shows that TBX18 upregulates FB-related genes in monolayer NRVMs model at d7 (upper panel), as well as in the Boyden chamber model (lower panel), which are reversed by A83-01, n= 4 biological replicates in each group, *$P$<0.05. In the Boyden chamber model (left), the top is transduced monolayer NRVMs, and the bottom is freshly isolated fibroblasts receiving 48h of starvation treatment. All multiple comparisons are performed with one-way ANOVA with the Bonferroni test.

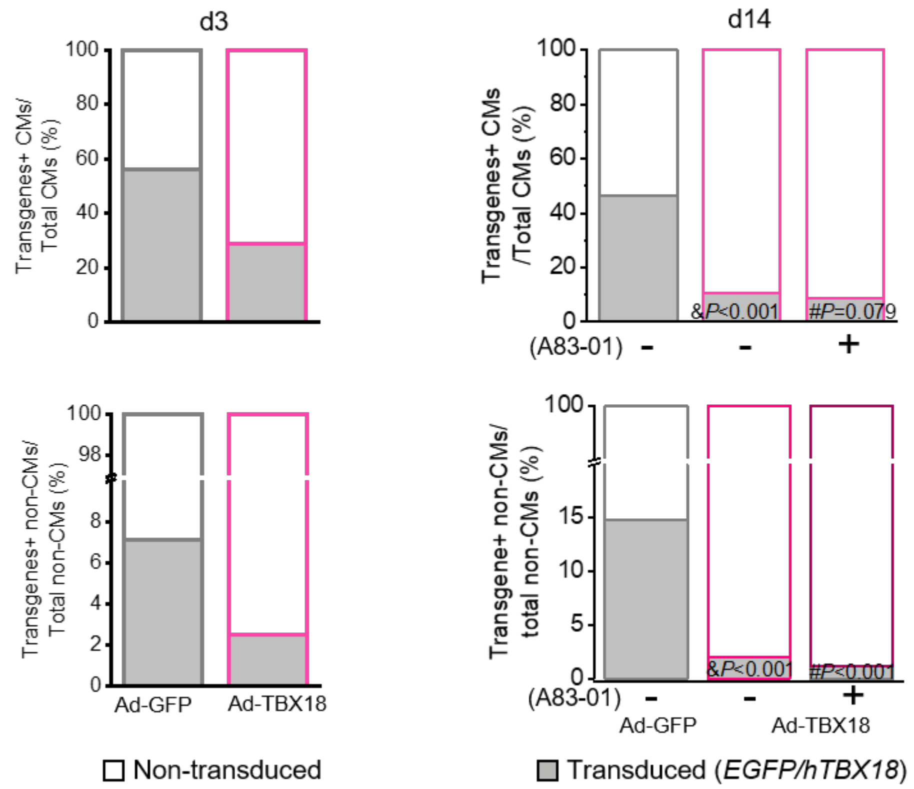


**Fig. S9. The proportion of *transgene*$^{+}$ CMs and nonmyocytes (non-CMs) at d3 and d14**. Most *transgene*$^{+}$ *(hTBX18 or EGFP)* cells are CMs, not non-CMs. The proportion of *hTBX18*$^{+}$ cells decrease from 28.6% to 10.5% in CMs and from 2.5% to 2.0% in non-CMs from d3 to d14, respectively. &$P$<0.001 compared to the ratio of *EGFP*$^{+}$ CMs (or non-myocytes), #$P$ compared to the ratio of *hTBX18*$^{+}$ CMs (or non-CMs), respectively, Chi-square test.

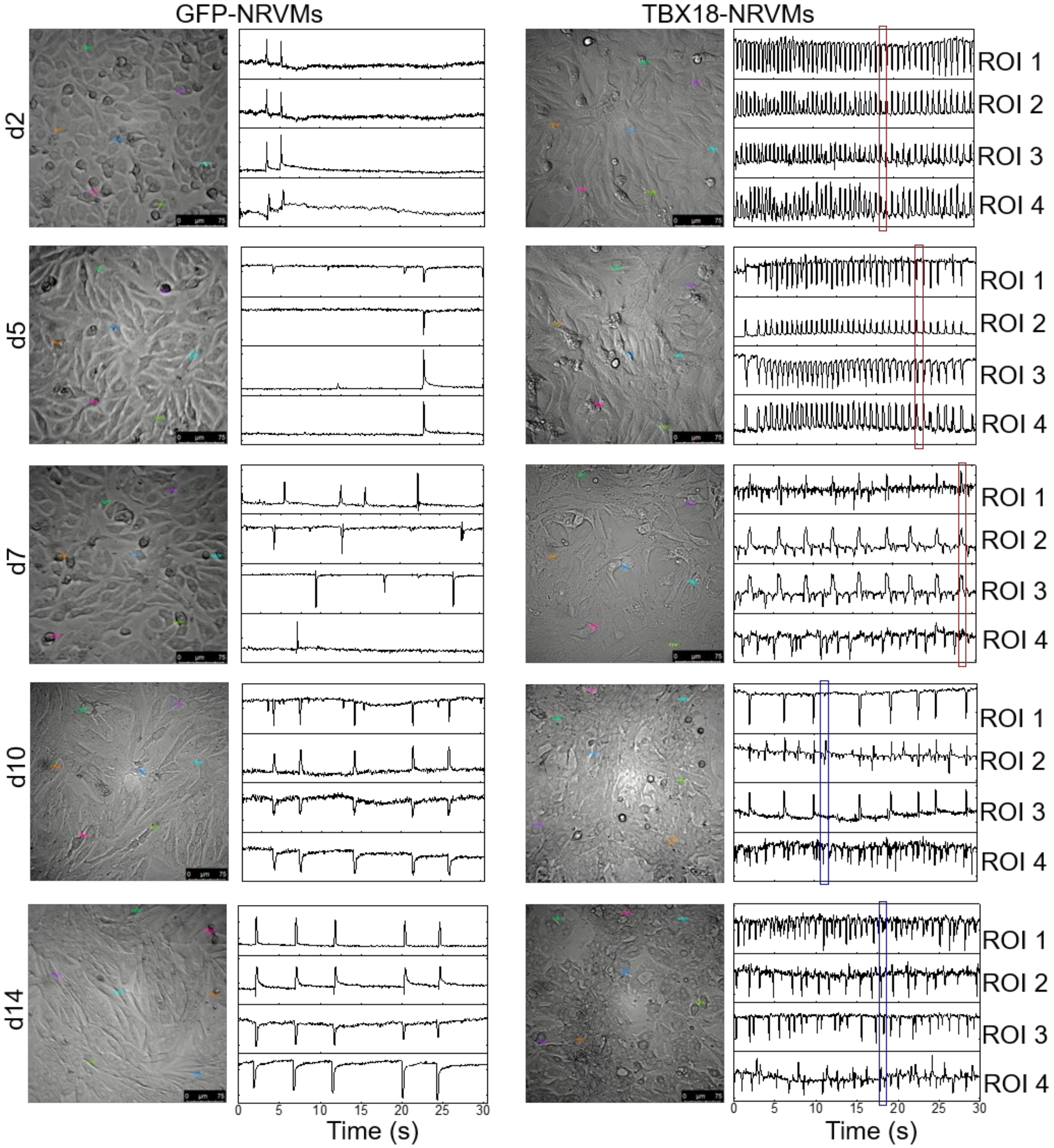


**Fig. S10. TBX18-NRVMs gradually acquire higher automaticity but lose synchrony.** The representative beat traces based on bright-field video indicate that TBX18-NRVMs acquire stronger automaticity but gradually lose synchronous contraction after one week. The red bar indicates syncytial beats among multiple regions of interest (ROIs), but the blue bar indicates asynchronous beats, 28 frames/s.

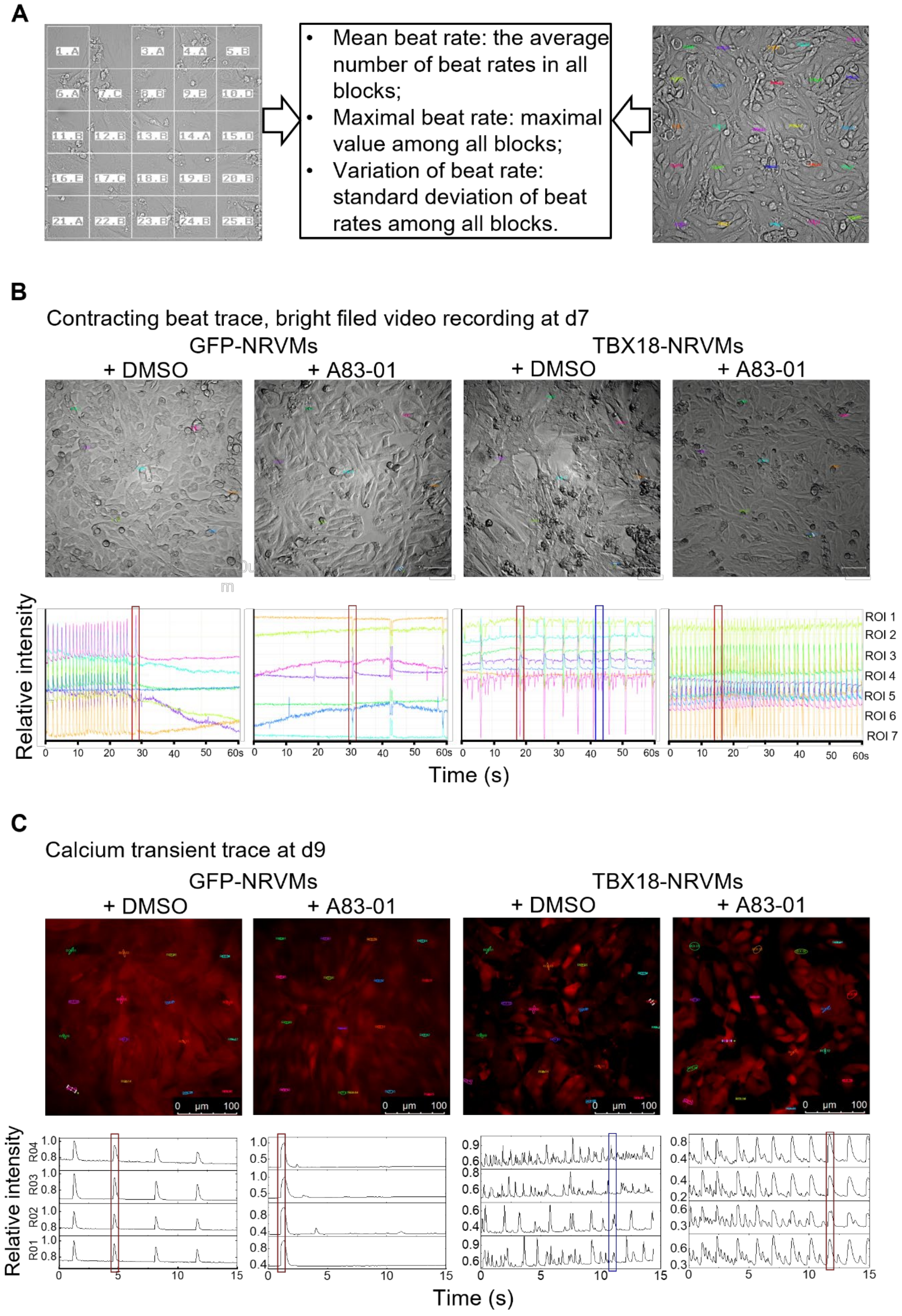
A
Mean beat rate: the average number of beat rates in all blocks;
Maximal beat rate: maximal value among all blocks;
Variation of beat rate: standard deviation of beat rates among all blocks.
B
Contracting beat trace, bright filed video recording at d7
GFP-NRVMs
TBX18-NRVMs
+ DMSO
+ A83-01
+ DMSO
+ A83-01
Relative intensity
ROI 1
ROI 2
ROI 3
ROI 4
ROI 5
ROI 6
ROI 7
Time (s)
C
Calcium transient trace at d9
GFP-NRVMs
TBX18-NRVMs
+ DMSO
+ A83-01
+ DMSO
+ A83-01
0 μm 100
Relative intensity
R04
R03
R02
R01
Time (s)

**Fig. S11. TBX18-NRVMs with A83-01 result in more synchronous beats compared to TBX18-NRVMs alone.** (**A**) The basic principle of video data analysis and the definition of critical indexes based on the pulse video analysis platform (https://pulsevideoanalysis.com). (**B, C**) Typical contracting beat traces show asynchronous beats of TBX18-NRVMs rescued by treatment with A83-01 (B, video recording under the bright field, 28 frames/s. C, calcium transient recording with Cal-590™ AM, 25 frames/s). The upper panel shows the screenshot of the video with multiple ROIs and the lower panel shows the beating trace of each ROI. The red and blue bar indicates syncytial and asynchronous beats among multiple ROIs, respectively.

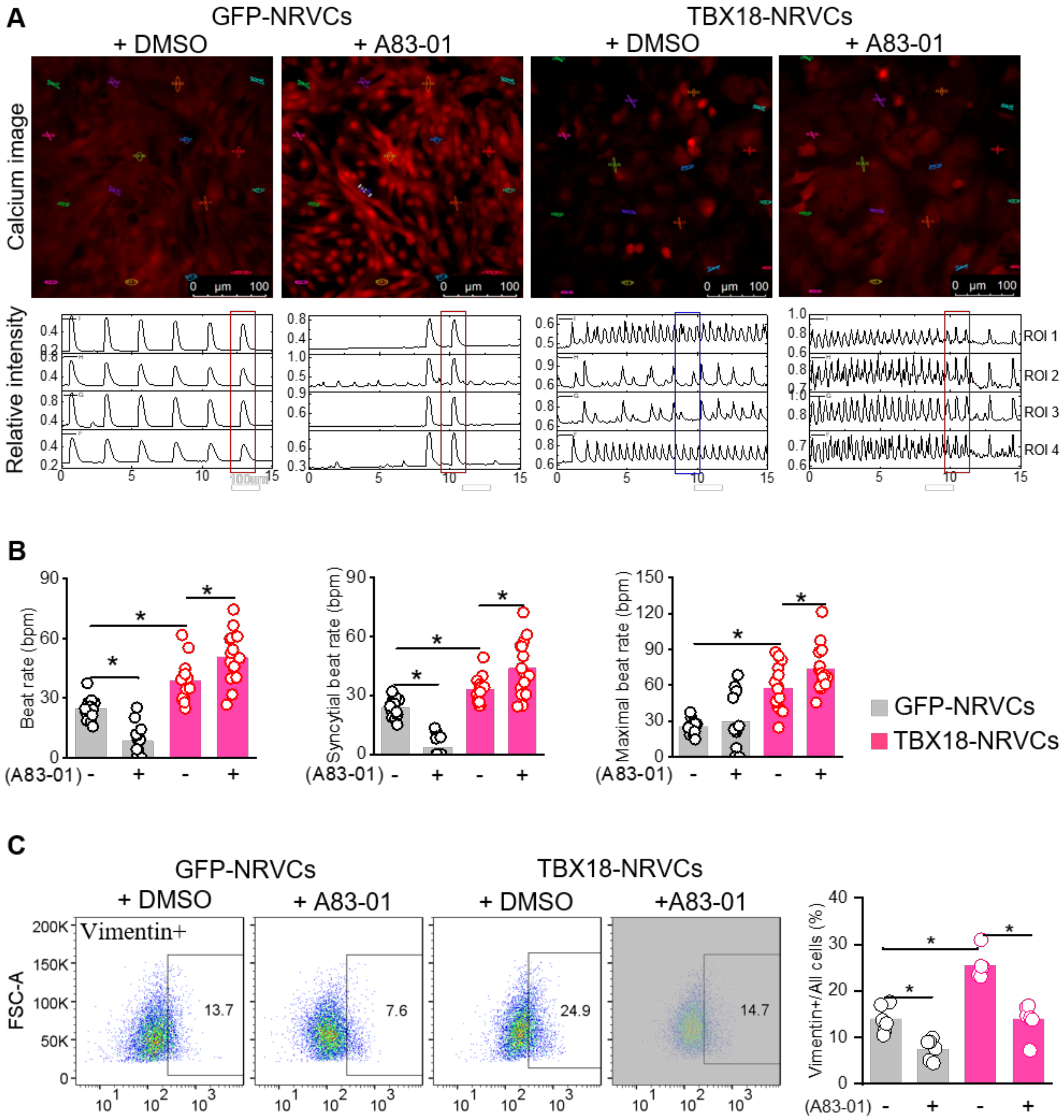


**Fig. S12. Treatment with A83-01 rescues asynchronous beats of TBX18-NRVCs at d8.** (**A**) Representative calcium transient traces show asynchronous beats of TBX18-NRVCs rescued by treatment with A83-01 at d8 (calcium transient recording with Cal-590™ AM, 25 frames/s). The upper panel shows the screenshot of the video with multiple ROIs and the lower panel shows the calcium transient trace of each ROI. The red and blue bar indicates syncytial and asynchronous calcium transient among multiple ROIs, respectively. (**B**) Measurements of spontaneous and syncytial contractions under bright field at d8 (n=15 biological replicates per group). **P*<0.05, one-way ANOVA with Bonferroni test. (**C**) Flow cytometry of NRVCs immunostaining against vimentin$^{+}$ cells at d8 after gene transfer. **P*<0.05, n=6 biological replicates in each group, one-way ANOVA with Bonferroni test.

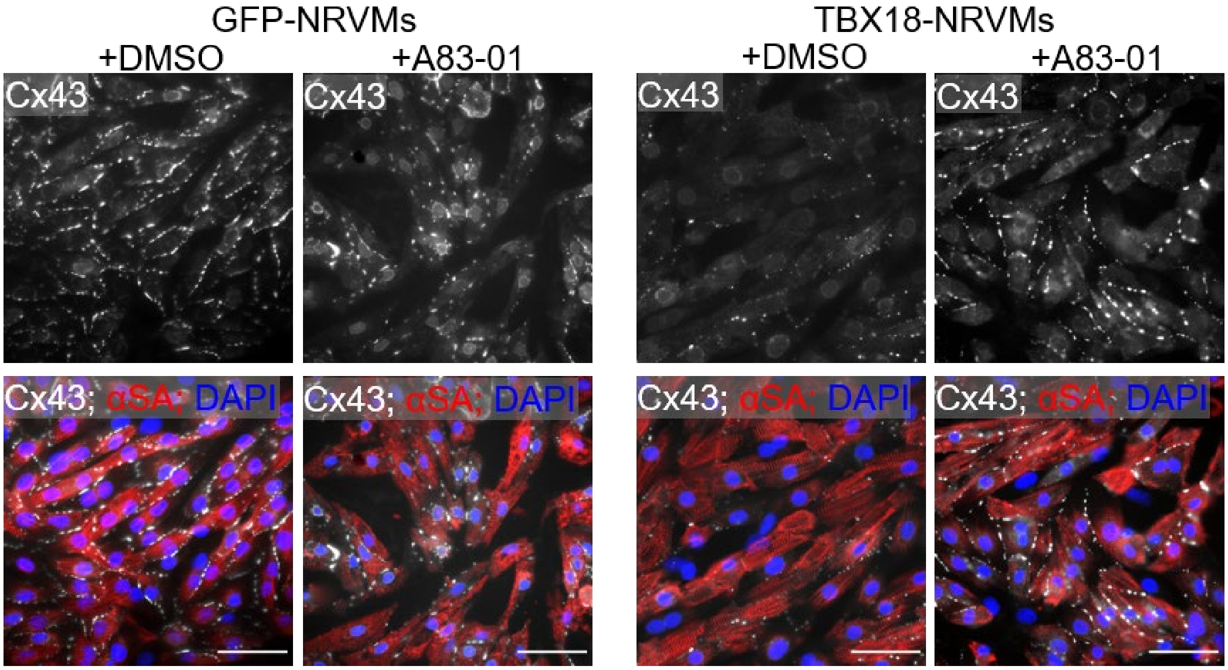


**Fig. S13. Inhibition of Tgfβ with A83-01 partially restores reduced connexin 43 by TBX18 in non-reprogrammed myocytes after 7 days.** TBX18-NRVMs display uniformly lower expression intensity of connexin 43 (Cx43), but treatment with A83-01 may preserve the expression levels of Cx43, especially for those non-reprogrammed CMs or non-transduced CMs, scale bar=50 µm.

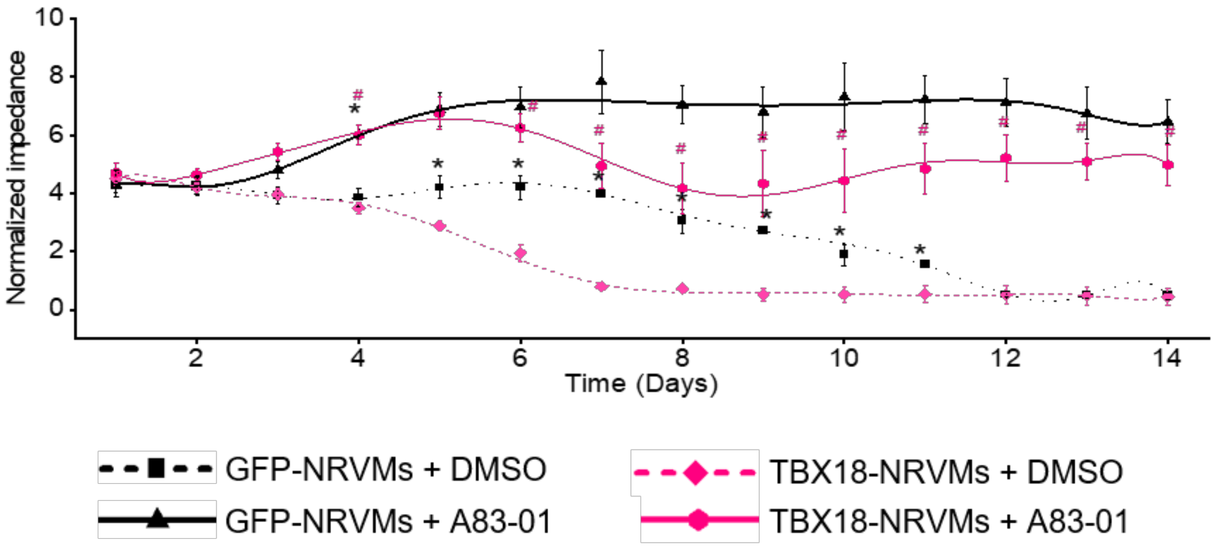


**Fig. S14. Tgfβ inhibition mitigates the drop in electrical impedance in TBX18-NRVMs for 2 weeks.** TBX18-NRVMs exhibit significantly lower impedance than GFP-NRVMs in the monolayer model, which are attenuated by its treatment with A83-01, n=8 biological samples in each group, *$P<0.001$ compared to TBX18-NRVMs+DMSO, #$P<0.001$ compared to TBX18-NRVMs+DMSO, one-way ANOVA with Bonferroni test.

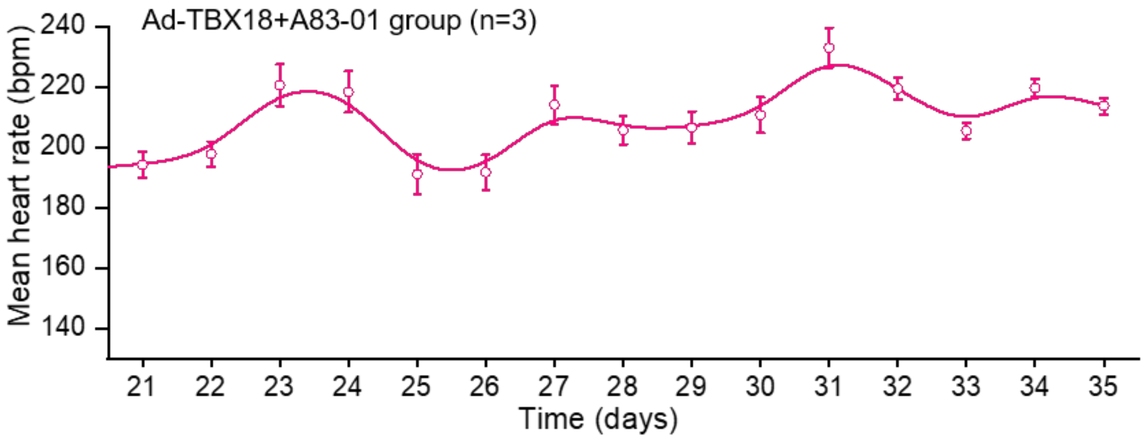


**Fig. S15. Animals with Adv-TBX18 plus A83-01 present persistently biological pacing after 5 weeks.** Three animals receiving Adv-TBX18 plus A83-01 treatment completed additional 2 weeks of ambulatory ECG monitoring after 3 weeks and presented persistent biological pacing with a higher mean heart rate.

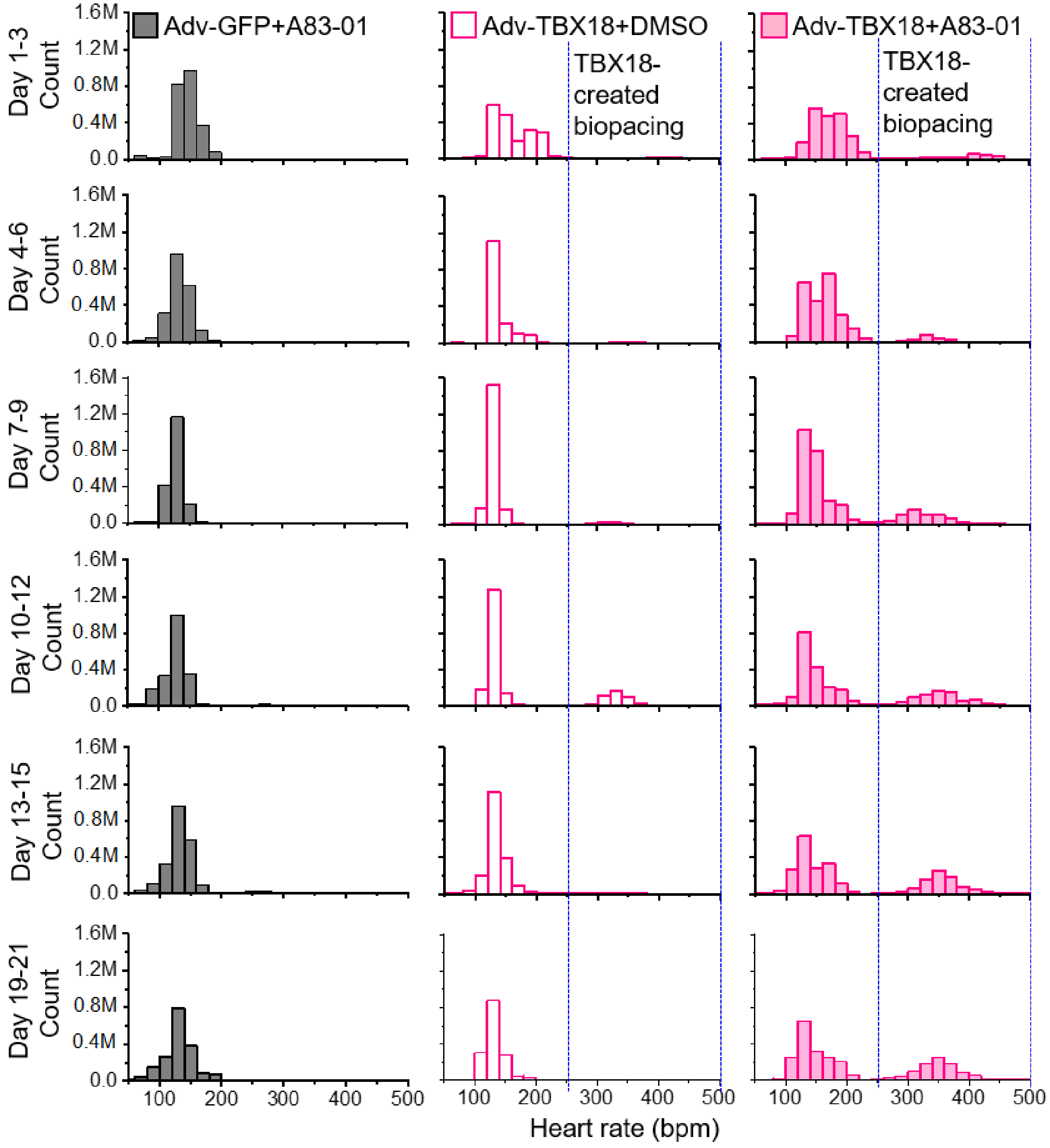


**Fig. S16. Temporal change of heart rate distribution among all ambulatory rats.** The average heart rates of every 4 beats were exported, and then the histogram was plotted with the heart rate on the X-axis and the count of heart rate on the Y-axis. Control animals present only one cluster of heart rates and unaltered junction rhythm with an average of 120-140bpm (left panel), and those rats receiving Adv-TBX18 with DMSO display an extra high frequency of heart rate (middle, between two blue lines) except for junction rhythm, about 300-400 bpm, especially at d10-d12, and then it wanes gradually. For animals with Adv-TBX18 and A83-01, the high-frequency area of heart rate (right, between two blue lines) does not shrink after 3 weeks. M, the abbreviation of million.

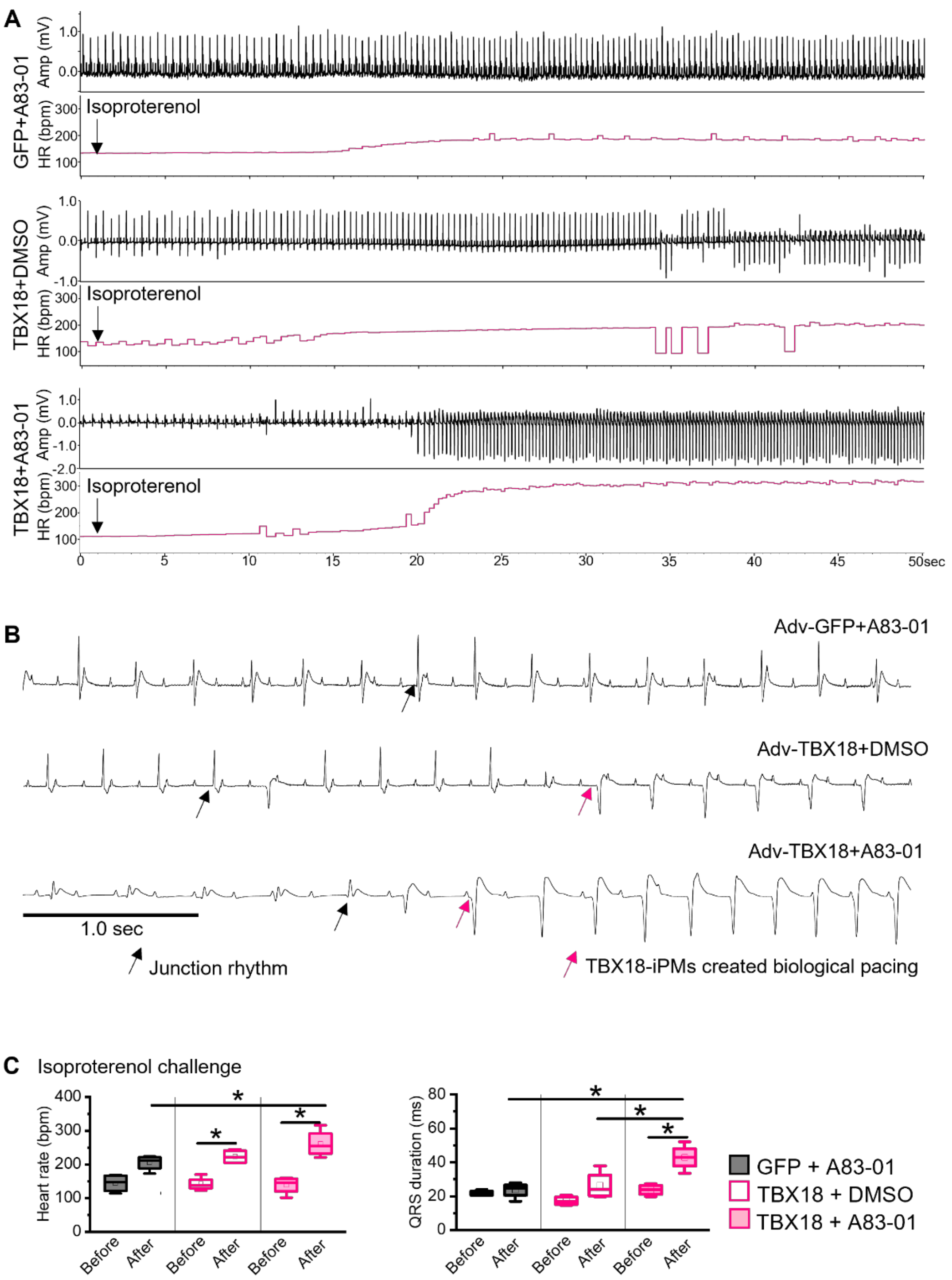


**Fig. 17. The response of TBX18-created biological pacing to the isoproterenol challenge test.** (**A**) Typical heart rate recording before and after the isoproterenol challenge. (**B**) Representative trace of surface ECG. Red arrows and black arrows indicate QRS complexes are from the apex and atrioventricular junction, respectively. (C) Heart rate analysis shows a significant increase in mean heart rate in animals with Adv-TBX18 plus A8301 after 10 minutes of isoproterenol challenge compared to the other 2 groups, as well as a wider QRS duration (n=4 rats; *, $P$<0.05), ANOVA with Bonferroni test.

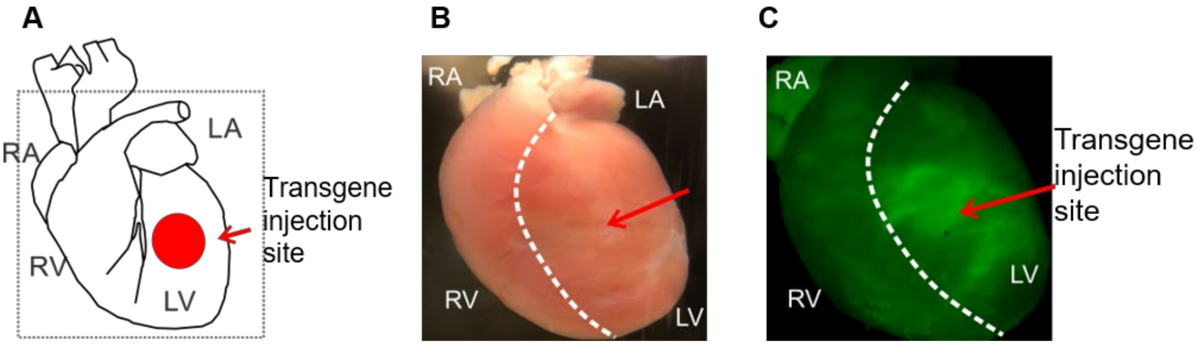


**Fig. S18. The schematic diagram of adenovirus injection at the left ventricle and the expression of reporter genes for optical mapping.** Adenovirus with transgene was injected at the anterior wall of the left ventricle (red circle, A), and then the expression of reporter genes (GFP or TBX18-Zsgreen) was detected under the fluorescent microscope to confirm the injection site.

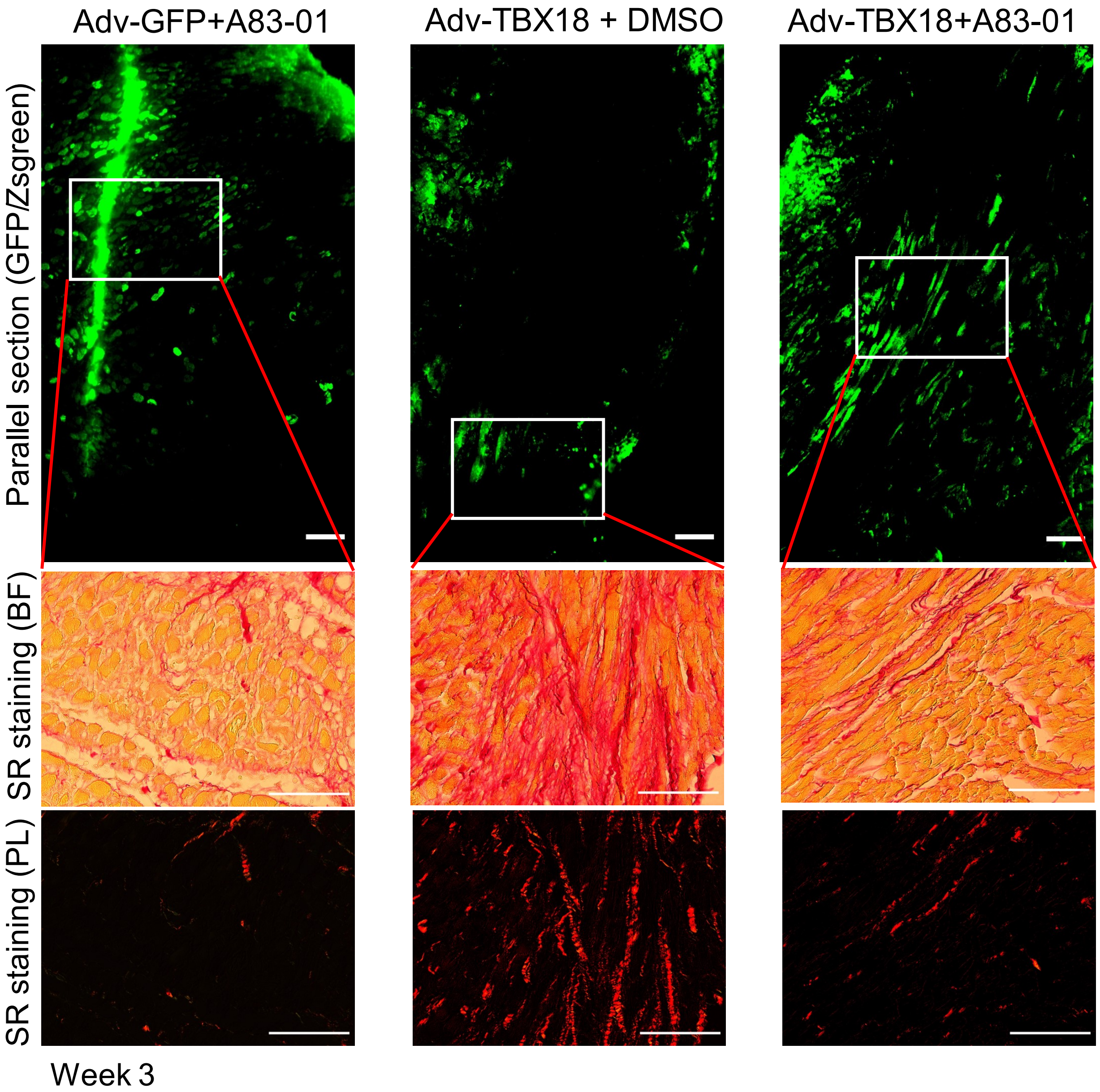


**Fig. S19. Fibrotic response at the TBX18-Zsgreen or GFP positive area at the edge of the injection site.** The top panel displays $GFP^+$ or TBX18-$Zsgreen^+$ cardiomyocytes at the edge of the injection site, and the white box indicates the corresponding area as below, scale bar=200 µm. The middle (bright field) and bottom (polarized light) panels show picrosirius red staining at the area of reporter $protein^+$ cells as above, scale bar=200 µm. The fibrotic response induced by TBX18 re-expression is remarkably attenuated when treated by A83-01.

**Legend of Movies**
**Downloading link for movies**
**https://1drv.ms/u/s!AgwEX2a0aSEFhCa-KM41Hk3lcKXd?e=zMfHSt**

**Movie S1. Typical spontaneous beating pattern and temporal changes of GFP-NRVMs.** Video recording under the bright field shows the beating pattern and temporal changes of GFP-NRVMs at d2, d5, d7, d10, and d14. GFP-NRVMs are mostly quiescent, occasional, and paroxysmal contractions, 28 frames/s.

**Movie S2. Typical spontaneous beating pattern and temporal changes of TBX18-NRVMs.** Video recording under the bright field shows the beating pattern and temporal changes of TBX18-NRVMs at d2, d5, d7, d10, and d14. TBX18-NRVMs show rhythmic beats within the first 5 days but begin to lose syncytial contractions. After one week, TBX18-NRVMs asynchronously present multiple spontaneous twitches and are incapable of generating syncytial contractions, 28 frames/s.

**Movie S3. TBX18-NRVMs with A83-01 result in more synchronous beats compared to TBX18-NRVMs alone** (bright field video recording, d7). Compared to GFP-NRVMs, TBX18-NRVMs at d7 exhibit multiple independently spontaneous twitches with a few syncytial contractions, but TBX18-NRVMs with A83-01 display synchronously rhythmic contraction, 28 frames/s.

**Movie S4. TBX18-NRVMs with A83-01 display more synchronous calcium transient compared to TBX18-NRVMs alone** (Calcium transient recording with Cal-590™ AM, d9**)**. TBX18-NRVMs at d9 show more active calcium transient but in an asynchronous manner, compared to GFP-NRVMs. In the contrast, TBX18-NRVMs with A83-01 display synchronous calcium transient propagation, 25 frames/s.

**Movie S5. Treatment with A83-01 rescues asynchronous beats of TBX18-NRVCs (Calcium transient recording, d8).** Unlike asynchronous calcium transient activity in TBX18-NRVCs at d8, TBX18-NRVCs with A83-01 present syncytial calcium contraction (Calcium transient recording with Cal-590™ AM, 25 frames/s).

**Movie S6. The activation pattern before and after atrioventricular node ablation in GFP-treated hearts.** The earliest activation site does not shift in the GFP-injected control hearts under sinus rhythm and upon atrioventricular node ablation.

**Movie S7. The earliest activation site shifts to the injection site before and after atrioventricular node ablation in TBX18-treated hearts**. After atrioventricular node ablation, the earliest electrical activation site shifted to the injection area in hearts with Adv-TBX18 in the presence of isoproterenol treatment.

**Movie S8. The earliest activation site shift to the injection site before and after atrioventricular node ablation in TBX18-treated hearts with A83-01**. Before and after atrioventricular node ablation, the earliest electrical activation site shift to the injection area only occurs in hearts with Adv-TBX18 plus A83-01 in the absence of isoproterenol treatment.

**Table S1. List of primers used in this study for real-time qRT-PCR.**

| Gene | Sequence |
|---|---|
| Rat Rpl4 QF | CACTTGGCGTAAGGCTGCTTCC |
| Rat Rpl4 QR | CCTGGCGGAGAATGGTGTTCCT |
| Rat Gjc1 QF | TCTGGCTCACTGTGCTGATTGT |
| Rat Gjc1 QR | GCTTCCAACGCATGGCATAGG |
| Rat Gja1 QF | GGATTGAAGAGCACGGCAAGGT |
| Rat Gja1 QR | ACACCAAGGACACCACCAGCAT |
| Rat Kcnj2 QF | GGCGGTGGATGCTGGTAATCTT |
| Rat Kcnj2 QR | GAACAGCCAGGAGAGCACGAAT |
| Rat Scn5a QF | CCATCCTGACCAACTGCGTGTT |
| Rat Scn5a QR | GGAAGGTGCGTAAGGCTGAGAC |
| Rat Hcn4 QF | TGAAGGCACCATCGGCAAGAAG |
| Rat Hcn4 QR | TGAGTAGAGGCGGCAGTAAGTATCC |
| Rat Gjc1 QF | TCTGGCTCACTGTGCTGATTGT |
| Rat Gjc1 QR | GCTTCCAACGCATGGCATAGG |
| Rat Thbs1 QF | GGCTTCATCTTCCTGGCTTCCT |
| Rat Thbs1 QR | ACTGACACCACTTGCTGCTTCC |
| Rat Col1a1 QF | TGTGCGATGGCGTGCTATGC |
| Rat Col1a1 QR | TCCTATGACTTCTGCGTCTGGTGAT |
| Rat Col3a1 QF | TTCTACACCTGCTCCTGTCATTCCT |
| Rat Col3a1 QR | CCATTCCTCCGACTCCAGACTTGA |
| Rat Eln QF | TGGTGCTACTGCTTGGTGGAGA |
| Rat Eln QR | CGTGGCTGCTGCTGTCTGATT |
| Rat Tgfb1 QF | GGACCGCAACAACGCAATCTATG |
| Rat Tgfb1 QR | TGCTCCACAGTTGACTTGAATCTCT |

| | |
|---|---|
| Rat tgfb2 QR | ATACACATCCAAGCAGGTCACCATT |
| Rat tgfbr1 QF | TAGCCAAGAGGAGATTATGAC |
| Rat tgfbr1 QR | GACAGCACAAGAGCGTAT |
| Rat tgfbr2 QF | TGTGAGAAGCCGCAGGAAGTC |
| Rat tgfbr2 QR | AGTGAAGCCGTGGTAGGTGAAC |
| Rat tgfbr3 QF | AATAGAAGACACGCTCCATATC |
| Rat tgfbr3 QR | GAGTCAATATAGCCTACCACAT |
| Rat tgfb3 QF | ATGATGATTCCTCCACACCGACTG |
| Rat tgfb3 QR | GCACTTACACGACTTCACCACCAT |

**Table S2. List of antibodies used for immunofluorescent staining, immunoblotting, and flow cytometry.**

| Primary antibody | Catalog | Manufacturer | Host | Ratio | Molecular weight |
|---|---|---|---|---|---|
| Western blot | | | | | |
| Anti-beta catenin antibody | ab16051 | Abcam | Rabbit | 1:600 | 95kDa |
| Anti-N-cadherin antibody | 33-3900 | Thermo Fisher | Mouse | 1:600 | 130kDa |
| Anti-vimentin antibody | ab24525 | Abcam | Chicken | 1:2000 | 50kDa |
| Anti-smad2/3 antibody | 8685S | CST | Rabbit | 1:1000 | 60/52kDa |
| Anti-phospho-Smad2 antibody | 3108S | CST | Rabbit | 1:400 | 60kDa |
| Anti-collagen I antibody | ab34710 | Abcam | Rabbit | 1:600 | 43kDa |
| Anti-connexin 43 antibody | C6219 | Sigma-Aldrich | Rabbit | 1:1000 | 43kDa |
| Anti-Hcn4 antibody | APC-052 | Alomone Labs | Rabbit | 1:200 | 130kDa |
| Anti-calnexin antibody | VPA00096 | Bio-Rad | Goat | 1:2000 | 75kDa |
| Immunostaining | | | | | |
| Anti-beta catenin antibody | ab16051 | Abcam | Rabbit | 1:400 | |
| Anti-N-cadherin antibody | 33-3900 | ThermoFisher | Mouse | 1:400 | |
| Anti-vimentin antibody | ab24525 | Abcam | Chicken | 1:600 | |
| Anti-actin, α-Smooth Muscle | A2547 | Sigma-Aldrich | Mouse | 1:300 | |
| Anti-actin, α-Smooth Muscle | ab32575 | Abcam | Rabbit | 1:400 | |
| Anti-collagen I antibody | ab34710 | Abcam | Rabbit | 1:200 | |
| Anti-connexin 43 antibody | C6219 | Sigma-Aldrich | Rabbit | 1:400 | |
| Anti-Hcn4 antibody | APC-052 | Alomone Labs | Rabbit | 1:100 | |
| Anti-Ki67 antibody | 550609 | BD Pharmingen | Mouse | 1:200 | |
| Anti-TBX18 antibody | SC-17869 | Santa Cruz | Goat | 1:200 | |
| Anti-α-actinin antibody | A7811 | Sigma-Aldrich | Mouse | 1:300 | |
| Flow cytometry | | | | | |

| | | | | |
|---|---|---|---|---|
| Anti-vimentin antibody | ab24525 | Abcam | Chicken | 1:600 |
| Anti-PDGFrα Antibody | ab203491 | Abcam | Rabbit | 1:400 |
| Anti-troponin I Antibody | A1-83509 | Scientific | Mouse | 1:400 |
| PE anti-CD31 Antibody | 555027 | Biosciences | mouse | 1:200 |
